# Integrating Secondary Structure Information Enhances Phylogenetic Signal in Mitochondrial Protein Coding Genes

**DOI:** 10.1101/2024.08.01.606191

**Authors:** Claudio Cucini, Francesco Nardi, Joan Pons

**Affiliations:** Department of Life Sciences, University of Siena. Via Aldo Moro 2, 53100 Siena, Italy; National Biodiversity Future Center (NBFC). Piazza Marina 61, 90133 Palermo, Italy; Departament de Biodiversitat Animal i Microbiana, Institut Mediterrani d’Estudis Avançats, IMEDEA (UIB-CSIC). Miquel Marqués 21, 07190 Esporles, Spain

**Keywords:** transmembrane regions, secondary structure, partitioning, mitochondrial genome, phylogenetics

## Abstract

Secondary structures have long been held to carry significant phylogenetic information, but the difficulty in collating primary sequence and structural information has frequently hindered its full application in phylogenetic studies. Here, we focus on the implementation of secondary structure information (inner, outer and transmembrane regions) to improve partitioning in datasets of mitochondrial Protein-Coding Genes for phylogenetic analysis. A new pipeline called TRAMPO was developed to partition alignment sites accordingly. The inclusion of this structural information in phylogenetic analyses was shown to produce an improvement in the phylogenetic signal of mitochondrial protein coding genes. We speculate that the observed, distinctive, compositional bias towards thymines in second codon sites of transmembrane regions, as well as selection, may be among the possible driving forces. This effect is especially relevant in lineages older than 50 Ma, where the signal of second codon positions becomes more relevant as first and third codon sites begin experiencing saturation.

## Introduction

In standard phylogenetic analyses, each individual position of an alignment is assumed to evolve independently of others, but one and the same evolutionary model (equilibrium frequencies, substitution matrix, rate of substitution) is applied across all positions (Felsenstein, 1978). It has, nevertheless, been repeatedly shown that the use of an inappropriate substitution model can have a detrimental impact on phylogenetic reconstructions, highlighting the necessity to carefully select the best-fitting model for a given sequence dataset (Posada and Crandall, 1998; Minin, 2003). Furthermore, since the initial implementation of DNA and protein sequences in molecular phylogenetics, researchers have realized that sequences of different origin (e.g. nuclear and mitochondrial) can display different equilibrium frequencies, a different substitution matrix and different rates of evolution, entailing that the use of a single evolutionary model is, *per se*, an unwarranted oversimplification. Such differences could be at play in Protein-Coding Genes (PCGs) across different codon positions (first, second and third codon sites), especially in the mitochondrial genome (Leavitt et al., 2013), as well as in genes encoding for ribosomal and transfer RNAs, where nucleotide positions in stems and loop regions evolve differently (Erpenbeck et al., 2007). Furthermore, several studies revealed that protein domains characterized by a different secondary structure are subjected to different evolutionary constraints (e.g. Le and Gascuel, 2010; Barbera and Cucini, 2022). Overall, this suggested that sequences should be analysed as different partitions, each evolving under an independent and specifically optimized evolutionary model, and that secondary structures should be accounted for in partitioned models (Pandey and Braun, 2020).

In modern phylogenetics, complex multigene sequence datasets, possibly including sequences from different origins and/or evolving under different selective constraints, are often split into smaller, uniform, partitions (e.g. PCGs are split by codon positions, ribosomal RNA by stems and loops). The best partitioning scheme is then identified that takes into account evolutionary heterogeneity without, at the same time, leading to an unwarranted increase in the number of model parameters (Kainer and Lanfear, 2015). Starting from a set of user-defined partitions that, based on current understanding, evolve under a reasonably uniform process, the analysis proceeds by iteratively merging smaller partitions into larger partition sets. Alternative partitioning schemes are compared, based on model fit as well as the number of free parameters, according to the Akaike Index Criterion (AIC) or Bayesian Index Criterion (BIC). This procedure can become computationally intensive as the numbers of starting partitions to merge increases, so the greedy heuristics were developed to this end (e.g. PartitionFinder; Lanfear et al., 2017).

This approach has been commonly implemented in phylogenetic analyses of datasets that include the full set of 13 mitochondrial PCGs, showing that partitioning of sequences by codon position (three partitions), or by codon position plus coding strand (six partitions), can lead to an increase in model fit and better phylogenetic insights than the analysis of the full dataset as a single partition (Pons et al., 2010; Bauzà-Ribot et al., 2012; Bover et al., 2019). These findings were further corroborated by studies describing the distinct evolutionary features of different codon positions and coding strands at different levels: nucleotide and amino acid composition, compositional variation among strands (AT- and GC-skews), among-site-rate variation, nucleotide substitution rate, phylogenetic content, selection (e.g. Pons et al., 2014).

Noteworthy, the procedure described above proceeds by merging smaller partitions into larger partitions, but does not split large partitions into smaller ones, even in the presence of model heterogeneity. As such, it is in the responsibility of the operator to identify a starting set of partitions based on categorization/discrimination rules (e.g. functional information such as by codon, by strand, etc.) that are deemed relevant. The proposed TRAMPO pipeline helps in creating a set of starting partitions, based on secondary structure information, to be evaluated in a partitioning analysis and eventually applied in the subsequent phylogenetic analysis.

The mitochondrion is an organelle of the eukaryotic cell able to produce energy via the oxidative phosphorylation (OXPHOS). From a structural standpoint, it is delimited by two double membranes, that in turn identify an inner space (matrix) as well as an intermembrane space and an outer space (cytoplasm). The mitochondrion has its own genetic material, independent of the nuclear genome, that in metazoans generally includes 37 genes - 13 PCGs, 22 tRNA, and 2 rRNA - *plus* a control region as the origin of transcription and replication (Boore, 1999). Each of the 13 encoded peptides is a multi-pass membrane protein that is embedded in the inner mitochondrial membrane. In fact, they are mostly composed of transmembrane (TM) helices spanning the inner membrane, but also include domains that protrude onto the mitochondrion matrix (MA) and the intermembrane space (IM) (Lloyd and McGeehan, 2013; Dennerlein and Rehling, 2015). Transmembrane helices can be predicted bioinformatically, based on primary sequence, using different types of software as: TMHMM (Möller et al., 2001), DeepTMHMM (Hallgren et al., in press), and HMMTOP (Tusnady and Simon, 2001). CCTOP (Dobson et al., 2015) was further developed to identify a consensus prediction based on different models. Each TM helix is composed of approximately 20 amino acids, whereas MA and IM regions can vary in length from as little as one amino acid to the more than 80 residues observed in the NAD5 protein. The fine scale structure of transmembrane regions (MA, TM, and IM regions) of human mitochondrial peptides was determined by Ian Logans (http://www.ianlogan.co.uk/TMRjpgs/tmrpage.htm).

In this study we analyse seven different datasets - multialignments of the 13 mitochondrial PCGs in three amphipod lineages (Crustacea), one hexapod lineage (Collembola=springtails) and three vertebrate sets (Vertebrata, Mammalia and Primates) - to investigate whether: a) different structural domains (MA, TM and IM) differ in their nucleotide and amino acid composition, as well as selection; and b) if this information can be deployed to improve evolutionary models in phylogenetic analyses. The seven groups were chosen to maximize phylogenetic representativeness, for their extension over different evolutionary time frames, and based on our experience with their systematics (e.g. Cucini et al. 2020; Jurado-Rivera et al., 2017). A new bioinformatic pipeline, named TRAMPO (TRAnsMembrane Protein Order), was created to classify nucleotide sites into the three structural classes (MA, TM, and IM). The effect of including structural information in partitioning and phylogenetic reconstructions was evaluated, and several sequence features, such as nucleotide and amino acid frequencies, A- and G-skew, Relative Synonymous Codon Usage (RSCU) and selection were specifically investigated across different domains.

## Material and Methods

### Datasets

Four datasets (complete sets of mitochondrial PCGs) were retrieved from previous studies: *Metacrangonyx* (Pons et al., 2019), *Pseudoniphargus* (Stokkan et al., 2018), *Hyalella* (Zapelloni et al., 2021) and springtails (Cucini et al., 2020) (Data S1). Three additional datasets (vertebrates, mammals and primates) were assembled from complete mitochondrial genomes available in GenBank in November 2021. One species per order, genus, and species was retained, respectively (Data S1). Altogether, the seven datasets were identified to have a broad taxonomic representation and a broad spectrum of evolutionary time-frames (25 Ma in *Hyalella* to 444 Ma in vertebrates; Zapelloni et al., 2021, Brazeau and Friedman, 2015).

### The TRAMPO (TRAnsMembrane Protein Order) pipeline

In order to partition mitochondrial PCGs into transmembrane helices (TM), mitochondrial matrix (MA) and intermembrane (IM) domains, a novel pipeline was created that is here named TRAMPO (pronounced trəmDpo). The pipeline is written in bash and python3 and includes some previously published code (Narakusumo et al., 2020; Cucini et al., 2021). TRAMPO operates in a dedicated conda environment (https://docs.conda.io/en/latest/) that is created upon setup and satisfies all its dependencies. TRAMPO is available, alongside installation and usage information, from a dedicated GitHub page (https://github.com/dbajpp0/TRAMPO). In brief, TRAMPO translates nucleotide sequences of interest into aminoacids, and then splits the alignment into transmembrane helices (TM), mitochondrial matrix (MA) and intermembrane (IM) domains by leveraging on an internal database of representative organisms whose domain structure was pre-determined.

The reference database was created starting from manually curated sets of mitochondrial PCGs available from Uniprot-SwissProt for representative species across major taxonomic ranks: *Homo sapiens* (vertebrates; NCBI accessions: YP_003024026-38), *Drosophila melanogaster* (pancrustaceans; YP_009047266-78), *Patiria pectinifera* (echinoderms; NP_008160-72), *Albinaria caerulea* (molluscs; NP_007329-41), *Lumbricus terrestris* (annelids; NP_008238-50), *Caenorhabditis elegans* (nematodes; NP_006953-64), and *Metridium senile* (cnidarians, NP_009252-53, NP_009255-65). TM, MA and IM domains were identified in each reference species using TMHMM (ver. 2.0; Möller et al., 2001) and HMMTOP (ver. 2.0; Tusnady and Simon, 2001) and a constrained consensus domain structure was obtained in CCTOP (ver. 1.1.0; Dobson et al., 2015). Domain structure was further curated in comparison with the high quality human model described by Ian Logan (http://www.ianlogan.co.uk/TMRjpgs/tmrpage.htm). Manual curation of domains generally included a) merging consecutive MA-TM-IM domains in *nad4* and *nad2* genes to reflect the larger, single domain observed in humans; b) revision of MA-IM order to ensure that all mitochondrial PCGs start with a MA domain, since this aspect is seldom predicted incorrectly (Möller et al., 2001).

For each new dataset, TRAMPO translates input nucleotide sequences into amino acids using the corresponding NCBI mitochondrial genetic code (invertebrate=5, vertebrate=2) and aligns the resulting amino acid sequences in MAFFT v7.397 under default parameters (Katoh & Standley, 2013). Then, TRAMPO splits PCG sequences into helices and domains based on the domain structure of the representative organism selected (invertebrates=dme, vertebrates=hsa). The domain structure is reported as a partition file in nexus format. In addition, TRAMPO produces tables, plots and statistics related to base and amino-acid composition, as well as codon usage. Focussing on functional differences, amino acids are aggregated in six functional groups (PAM250 matrix, Dayhoff et al., 1978): G1 (Small, A, G, P, S, T); G2 (Acid and Amide D, E, N, Q); G3 (Basic H, K, R); G4 (Hydrophobic I, L, M, V); G5 (Aromatic F, W, Y); G6 (Sulphur polymerization, C). TRAMPO further outputs several partitioning schemes at the nucleotide level (by codon, by strand, by domain, as well as combinations thereof) to facilitate subsequent PartitionFinder analyses.

### Domain analyses

As discussed above, numerous studies have shown that splitting mitochondrial PCGs by codon position (three charsets: first, second, and third codon sites) and strand (two charsets: plus and minus), for a total of six partitions, significantly improves model fit and leads to improved phylogenetic reconstructions. In this study, we evaluated whether additional partitioning by secondary structure domain (three charsets: matrix-associated (MA), transmembrane (TM), and intermembrane (IM)) can lead to an additional improvement. The initial partitioning schemes evaluated were: (i) all sites as a single partition; (ii) partitioning by codon (three charsets); (iii) partitioning by codon and strand (six charsets); (iv) partitioning by codon and domain (nine charsets); and (v) partitioning by codon, strand, and domain (18 charsets). Based on initial results, an additional partition scheme was implemented where MA and IM domains are merged into a single set (referred to as MA+IM). This resulted in a new partition scheme (vi) by codon, domain (TM vs MA+IM), and strand (12 charsets).

These six partition schemes were processed (i.e. merged/optimized) using the PartitionFinder algorithm (Lanfear et al., 2017) implemented in IQ-TREE2 v2.07 (Minh et al. 2020). The rcluster algorithm was used to merge partitions, in conjunction with the AICc criterion, to evaluate model fit along merging iterations (--merge rcluster --rcluster 100 --rcluster-max 1000 --merit AICc). ModelFinder (Kalyaanamoorthy et al., 2017), as implemented in IQ-TREE2, was used to estimate the best nucleotide substitution model for each partition, and IQ-TREE2 to build Maximum Likelihood trees. Branch lengths across partitions were independently estimated (option -Q in IQ- TREE2) in order to study the phylogenetic signal in each partition. To speed up the analysis, only three nucleotide substitution models were evaluated (option --mset HKY, TN, GTR). This is justified because unequal frequency models always outperform equal frequency models in AT-rich datasets, such as mitochondrial genomes. Besides, gamma (G) and invariant (I) models were used to model among site rate variation (option -mrate G,I), while the invariant *plus* gamma (I+G) model was not included as the two parameters are inextricably linked (Sullivan et al., 1999). Considering that the number of parameters was different in different models, partitioning schemes and models were compared based on corrected Akaike Information Criterion (AICc). BIC (Bayesian Information Criterion), in turn, was not applied as the number of species varied extensively, particularly in the vertebrate datasets.

In order to investigate the possibility that different partitioning schemes support different phylogenetic trees, the approximately unbiased test (AU; Shimodaira, 2002) was used to identify statistically supported incongruences among tree topologies. The AU test, as implemented in IQ- TREE2, was conducted with 10 thousand replicates by comparing each tree with the best tree (i.e., the tree obtained under the best partitioning scheme). The tree space was further explored in TreeDist (Smith, 2022) using both clustering information based on K-means and quartet distances, as they are more sensitive to evolutionary relationships (i.e. tree topology) than to marginal features like tree shape (Smith, 2022). Mapping of distances using principal coordinates analysis (PCoA) was preferred over Sammon and Kruskal mappings since the former generally replicates the original tree-to-tree distances more accurately (Smith, 2022).

Tree topologies estimated in IQ-TREE2 were midpoint-rooted in FigTree v1.4.4 (https://github.com/rambaut/figtree) to perform selection analyses. The possibility of ongoing selection in TM and MA+IM regions was assessed in HyPhy v2.4.5.21hf (https://www.hyphy.org/; Pond et al., 2020) using the MEME (Mixed Effects Model of Evolution; Murrell et al., 2012) and the FEL (Fixed Efects Likelihood; Pond and Frost, 2005) methods. The former (MEME) assesses whether individual sites are under episodic positive or diversifying selection (i.e. detect sites evolving under positive selection under a proportion of branches). The second (FEL), in turn, detects sites under pervasive positive diversifying selection or sites under negative selection at *p* ≤ 0.1. The possibility of positive selection acting at the level of specific biophysical and biochemical amino-acid properties was further investigated using TreeSAAP (Woolley et al., 2003). This analysis compares changes in amino acid chemical properties at non-synonymous sites across a fixed phylogeny and identifies sites of each class under positive selection. Sequences were analysed as a single concatenated protein and sites were subsequently assigned to different domains (MA, TM, IM) based on partitions obtained with TRAMPO. Values above 3.090 in categories 6, 7 and 8 were considered significant since amino acid changes display the strongest statistical support (*p* < 0.001). A pairwise Chi-square test was performed to assess if the number of amino acids under selection in TM helices was different from the number of sites under selection in MA+IM domains. Chi-square tests were performed with a 2×2 contingency table using the GraphPad online resource (https://www.graphpad.com/quickcalcs/contingency1).

## Results

Based on TRAMPO domain-based partitioning, aligned positions were assigned to the same domains (MA, TM, and IM) predicted in *H. sapiens* (vertebrates, mammals and primates) and *D. melanogaster* (Collembola, *Hyalella*, *Metacrangonyx* and *Pseudoniphargus*) in all the seven datasets. Final alignments and domain designation for the seven datasets are provided as Supplementary Material (Data S1). As expected, the number of positions in transmembrane helices (TM) outnumbered those in matrix and inner membrane domains (MA, IM; Table 1) in all datasets and genes, except in the short *atp8* and *cox2* sequences that include only one and two transmembrane helices, respectively.

**Table 1.**
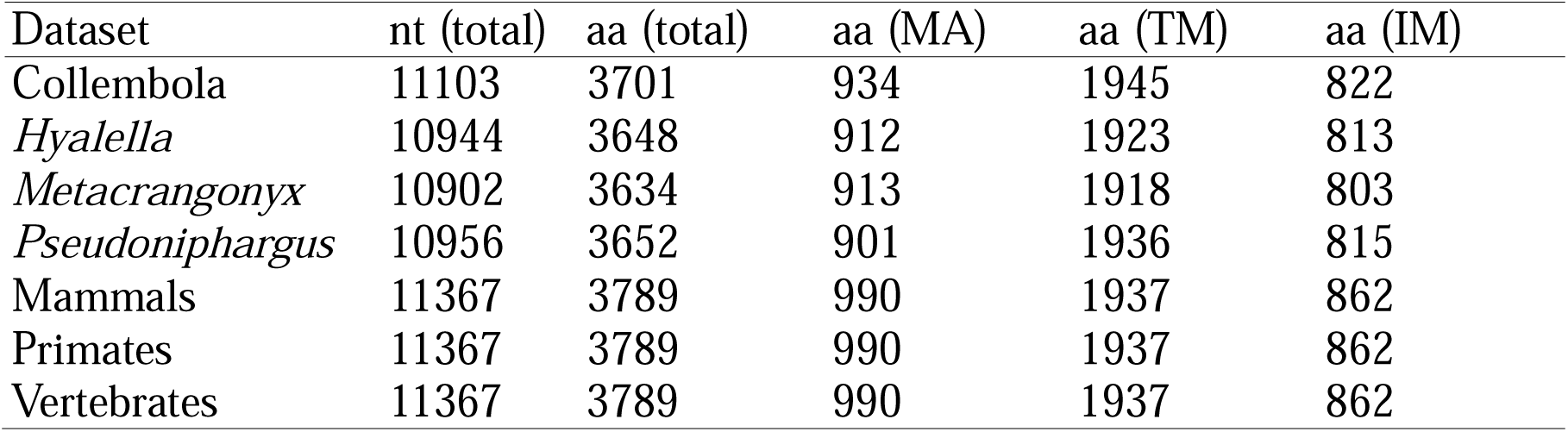
Length of the concatenated sequences of the 13 mitochondrial protein coding genes for the seven datasets, at the nucleotide (nt) and amino acid (aa) level, as well as the number of amino-acids assigned to each domain type (MA, TM, IM).

| Dataset | nt (total) | aa (total) | aa (MA) | aa (TM) | aa (IM) |
| --- | --- | --- | --- | --- | --- |
| Collembola | 11103 | 3701 | 934 | 1945 | 822 |
| *Hyalella* | 10944 | 3648 | 912 | 1923 | 813 |
| *Metacrangonyx* | 10902 | 3634 | 913 | 1918 | 803 |
| *Pseudoniphargus* | 10956 | 3652 | 901 | 1936 | 815 |
| Mammals | 11367 | 3789 | 990 | 1937 | 862 |
| Primates | 11367 | 3789 | 990 | 1937 | 862 |
| Vertebrates | 11367 | 3789 | 990 | 1937 | 862 |

Nucleotide composition, as well as A- and G-skews, were observed to differ both across codon positions (first, second, and third) and coding strands (positive and negative; Fig. 1, and Fig. S1). As expected, third codon sites, irrespective of coding strand and domain type, showed the highest A+T content relative to first and second codon sites (Fig. S2). This was more pronounced in the four pancrustacean datasets (springtails and amphipods), notably A+T rich, compared to the three vertebrate datasets (Fig. S2). Common to all datasets, the most striking result was that second codon sites of TM helices exhibited a very distinct base composition pattern compared to second codon sites of MA and IM domains, as well as from other partitions. These sites (i.e. second codon positions of TM helices) showed extremely low A-skew values (∼0.2), indicating an intra-strand excess of Ts relative to As (Fig. 1 and Fig. S1). In the four pancrustacean datasets an additional, less pronounced, difference was observed between second codon positions of TM helices encoded in the two strands (Fig. 1a,b and Fig. S1). In one dataset - a possible idiosyncrasy of springtails - second codon sites on the negative strand of MA regions displayed an even more extreme A-skew (< 0.2). In some lineages, first and third codon sites of TM regions were also retrieved as independent clusters (see Fig. 1 and Fig. S1). At variance with second and third codon sites, first codon sites exhibited a more equilibrated base composition (Fig S1, Fig S2). In summary, the key result concerning base composition was that second codon sites of TM helices stand out for their T richness relative to other partitions.

**Figure 1.**
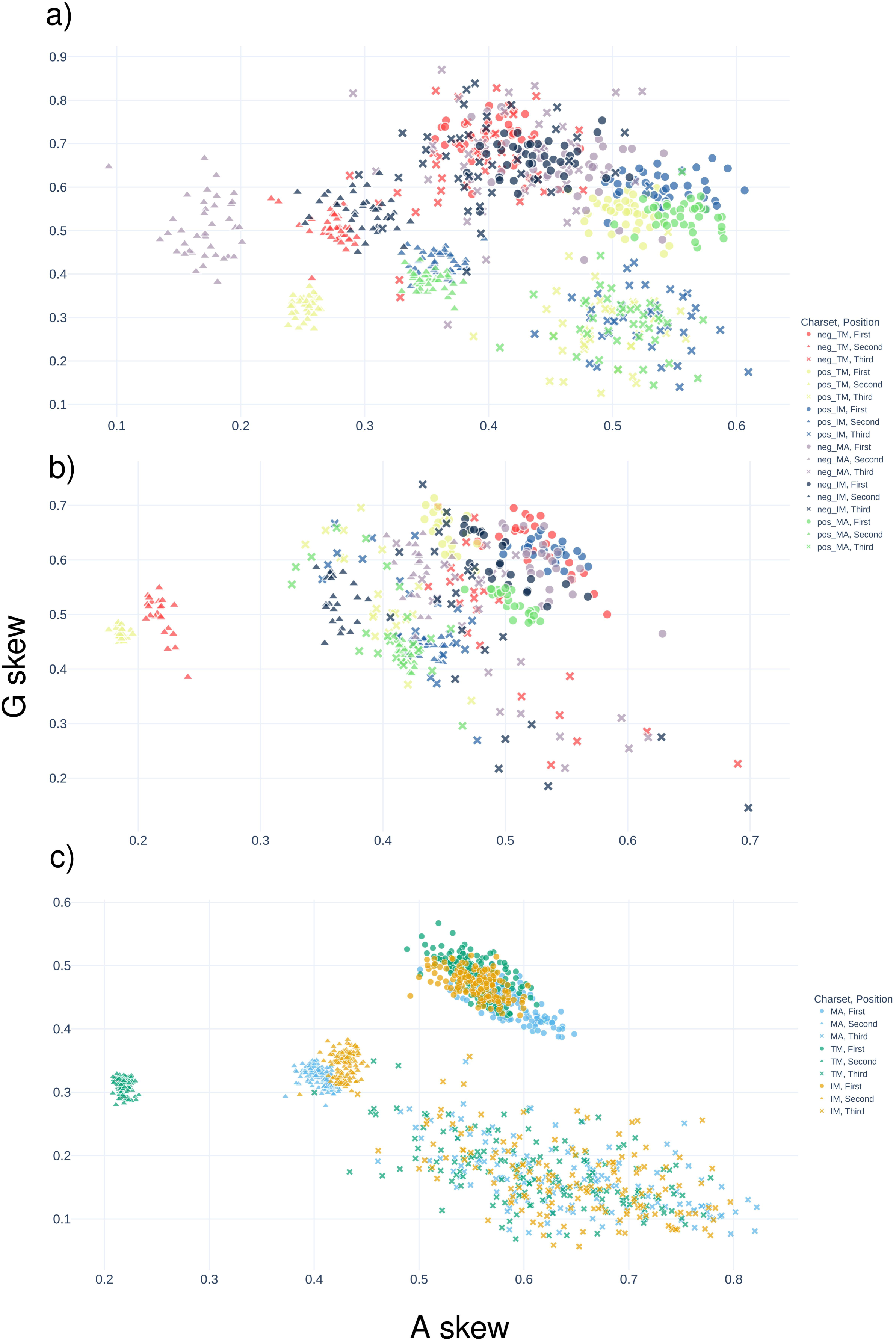
Scatter plots of A-skew (x-axis) vs G-skew (y-axis) for different domains (IM, MA, TM), strand (positive and negative) and codon position for selected datasets: Collembola (a) *Metacrangonyx* (b) and Vertebrates (c). Partitions are named as ‘strand_domain_codon position’. See Supplementary Figure S1 for other datasets. Due to the presence of only one gene encoded in the negative strand of vertebrates, panel (c) does not display strand differences.

TM helices also showed a different amino acid frequency relative to MA and IM domains in all the seven datasets (Fig. 2 and in Fig. S3). Overall, the frequency of amino acids in functional groups G2 (Acid and Amide) and G3 (Basic) were lower in TM helices compared to MA and IM domains, with marginal differences in G1 (Small) group. On the other hand, frequencies of amino acids in group G4 (Hydrophobic) were higher in TM helices, with marginal differences in group G5 (Aromatic). The only exception, concerning group G4, was observed in the negative strand of the MA regions in the springtail dataset (Fig. 2a), the same partition where lower A-skew frequencies were observed at second codon sites (see above). Differences among strands were of lesser extent than between transmembrane helices and matrix/intermembrane domains (Fig. 2a,b). Finally, at the individual codon level, RSCU analysis showed that AT-rich codons are generally preferred over GC-rich codons, with minor differences among the three regions. The three vertebrate datasets showed some additional differences relative to the pancrustacean datasets, in the proportion of Leucine and Serine codons (data not shown).

**Figure 2.**
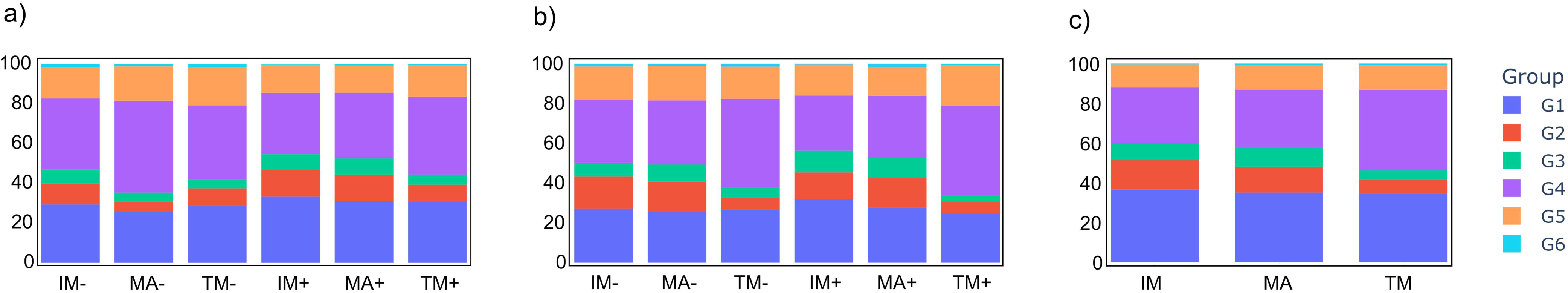
Frequency of amino-acids in different chemical classes in different domains (IM, MA and TM) and strands (positive and negative) for three selected datasets: Collembola (a), *Metacrangonyx* (b), and Vertebrates (c). Partitions are named as ‘strand_domain_codon position’. See Supplementary Figure S3 for other datasets. Due to the presence of only one gene encoded in the negative chain of vertebrates, panel (c) does not display strand differences.

The best partitioning schemes, following PartitionFinder merging and optimization, were obtained starting with 18 (TM/MA/IM *plus* by strand *plus* by codon; in *Pseudoniphargus*, *Metacangronyx*, vertebrates and primates) or 12 (TM/MA+IM *plus* by strand *plus* by codon; in Collembola, *Hyalella* and mammals) partitions (Fig. 3, Table S1). Noticeably, initial schemes that resulted in the best optimized schemes invariantly included transmembrane sites as a separate starting partition (i.e. TM/MA/IM in 18 starting partitions or TM/MA+IM in 12 starting partitions). Using 18 (TM/MA/IM *plus* by strand *plus* by codon) or 12 (TM/MA+IM *plus* by strand *plus* by codon) starting partitions, in turn, led to similar AICc values, suggesting that the key difference is between TM and MA+IM, and not between MA and IM.

**Figure 3.**
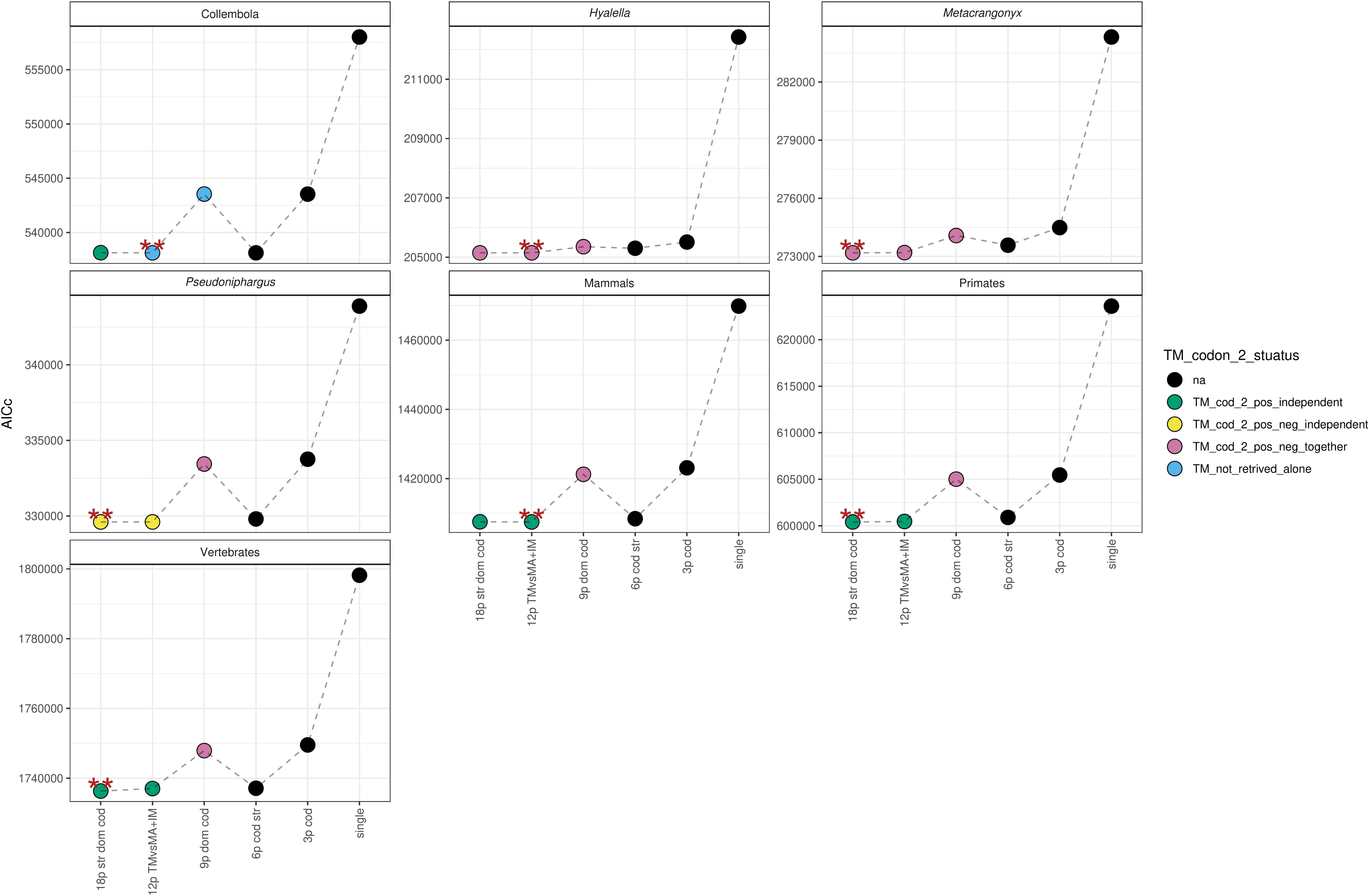
Quality of the partitioning schemes obtained after Partition Finder merging/optimization starting from six different partitioning schemes (x axis). AICc values are reported on the y-axis. Colour coded is the status of second codon positions of the positive strand in TM helices following PartitionFinder merging/optimization. Double asterisk identifies the partitioning scheme with lowest AICc in each dataset.

The typical use of partitioning by codon and strand (6 charset) performed well in many lineages but produced slightly higher AICc than in analyses with 18 and 12 starting blocks. On the other hand, partitioning by codon only performed poorly, as well as splitting codons further by transmembrane helices but not by strand (Fig. 3, Table S1). As expected, no partitioning resulted in the worst results.

A global look at the final partitions (merged and optimized with PartitionFinder) obtained from the 18 (TM/MA/IM *plus* by strand *plus* by codon) and 12 (TM/MA+IM *plus* by strand *plus* by codon) starting partitions in the seven lineages showed some transversal features (Table S2). TM helices were consistently separated from MA+IM domains at the second codon positions (Fig. 3, Tab. S2), except in Collembola and *Hyalella*, where this is true for the positive strand only (limited to the three partitioning schemes that include domain information). In the three vertebrate lineages, this separation is only observed on the positive strand when strand metrics were included. This aligns with the observation that vertebrates have only one gene present on the negative strand, and therefore starting charsets are significantly reduced in size and signal. In amphipod lineages, the second codon positions of TM helices were clustered into different sets. They were further split by strand into neg_cod2_TM and pos_cod2_TM, as seen in *Pseudoniphargus*, or merged into a single partition (cod2_TM) as in *Hyalella* and *Metacrangonyx*. Collembola results varied depending on the partition scheme implemented (Table S2). Altogether, this shows that TM helices and MA+IM domains are optimally placed in different partitions. This is in line with the observation that second codon sites of TM helices display a unique nucleotide and amino acid composition, thus justifying the use of a different evolutionary model (i.e. a different partition). In practical terms, this entails that starting partitioning schemes for PartitionFinder should also include partitions based on secondary structure, either TM/MA/IM, as in the 18 partitions scheme above, or minimally TM/MA+IM, as in the 12 partitions scheme above. Additional trends were observed concerning first and third codon positions: a) first codon positions in the negative strand were retrieved as a separate partition or associated with second codon positions, as in the three vertebrate datasets where only one gene is present in the negative strand; b) third codon positions on the negative strand are retrieved as a separate partition; c) first and third codon positions in the positive strand are retrieved as a separate partition or further split in TM helices and MA+IM domains, with contrasting patterns in Amphipoda, on one end and Collembola *plus* the three vertebrate datasets on the other.

Despite the extensive differences among the six partitioning schemes tested, and associated differences in the resulting trees, none of the topologies were incongruent based on AU-test relative to that obtained from merging 18 partitions. This indicates that the observed differences among phylogenetic trees are generally small, or receive little support. A closer look at the tree space using PCoA based on quartet distances and clustering constructed using K-means (Fig. 4) resulted in silhouette coefficients lower than 0.5, except for Collembola, indicating that the clustering structure resulted in a suboptimal visualization of the composition of the primary tree spaces. The tree space for *Hyalella* and *Pseudoniphargus* datasets were not analysed, since all partitioning schemes resulted in the same tree topologies. Tree distances relative to focal tree (i.e. obtained with the 18 partitioning scheme) were small in most datasets (i.e. yellow-green-orange lines in Fig. 4), except for *Metacrangonyx* and vertebrates. The analyses of those distances show that trees are mostly grouped into two or three clusters, except for vertebrates, where trees appear to be more spread apart in the tree space.

**Figure 4.**
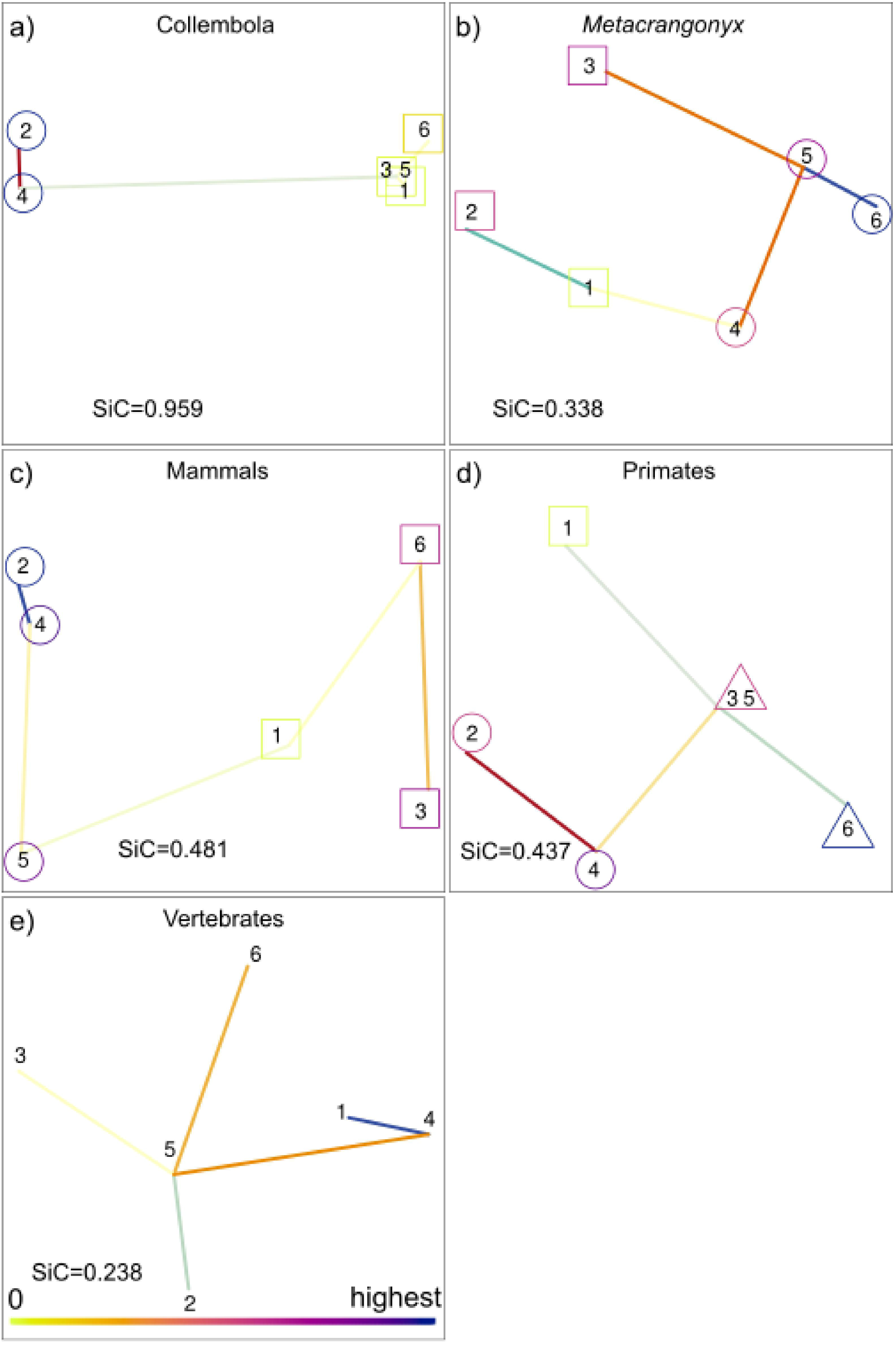
Principal component analyses of tree space of tree topologies obtained using different partition schemes. Tree numbers correspond to partition schemes in Fig. 3. Tree clusters based on quartet distances and K-means are indicated with different symbols. Colour bar in panel e) denotes tree distances from focal tree (i.e. tree from partition scheme obtained by merging/optimization of 18 partitions).

The tree obtained by merging the 18 starting partitions in each dataset (i.e. almost invariantly the best scheme overall, Fig 3) was also used to study selection. TM helices were explicitly compared with MA+IM domains, as all previous analyses indicated that a major difference exists, in terms of equilibrium frequencies as well as evolutionary model, between TM and MA+IM, whereas the difference between MA and IM appears to be less pronounced. The presence of episodic positive or diversifying selection (i.e. positive selection) was explored using the MEME test in HyPhy. As expected, few amino acid positions were under positive selection (*p* ≤ 0.1, Fig. 5a). Their frequency, in TM helices, ranged from ∼1-2% in amphipods to ∼3-5% in mammals, vertebrates, springtails and primates. The Chi-squared test rejected the null hypothesis (*p* ≤ 0.05) of equal distribution of sites under positive selection among TM helices and MA+IM domains in *Hyalella*, mammals and vertebrates (Fig. 5a), indicating that in these datasets positive selection is more frequent in TM helices compared to MA+IM domains. The FEL test (*p* ≤ 0.1) detected a generally lower frequency of sites under pervasive positive diversifying selection (∼0.05-0.35% in TM helices, Fig. 5b) with no significant differences between TM and MA+IM domains. On the other hand, it showed that most sites are under negative selection (∼75-95% in TM helices, Fig. 5c) with a statistically significant difference between TM and MA+IM regions in *Hyalella* and *Metacrangonyx* (where negative selection prevails in MA+IM domains) and vertebrates (where negative selection prevails in TM helices; Fig. 1c).

**Figure 5.**
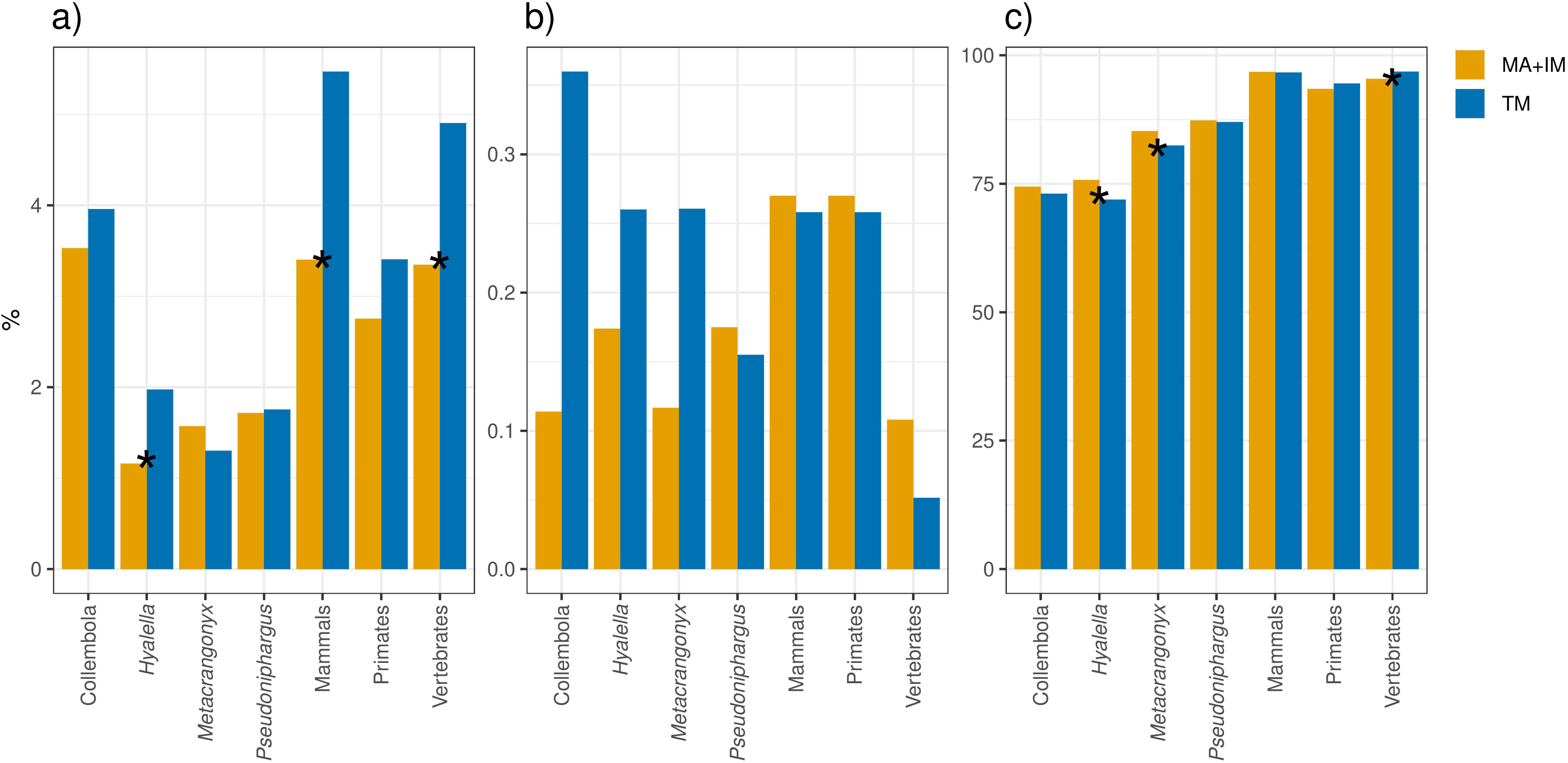
Percentage of amino-acid sites under selection in TM vs. MA+IM domains as predicted (a) with MEME (episodic diversifying positive selection), (b) with FEL (pervasive positive diversifying selection), (c) with FEL (negative selection). Asterisks indicate significant differences between TM and MA+IM regions (*p* ≤ 0.05).

Eleven individual physicochemical properties were found to be under selection in mitochondrial PCGs (Table 2). Nonetheless, few were consistently identified across the seven datasets under study: equilibrium constant ionization of COOH (EC) and solvent accessible reduction ratio (SA) were identified in all datasets; alpha-helical tendencies (AH) and power to be at the C-terminal (PC) in six out of seven datasets; long-range non-bonded energy (LR) in five. From 4 (*Hyalella*) to 10 (Primates) biochemical properties were hypothesized, overall, to be under selection in different groups. Interestingly, the power to be at the middle of alpha-helix, relevant for the structural categorization proposed, was identified as under selection in the three vertebrates datasets but in none of the invertebrate datasets. The number of amino acid sites identified as under selection (Fig. 6), limited to traits that were indeed identified as under selection (Table 2), varied widely, ranging from 1 to 14 sites in TM helices depending on taxon and trait (Fig. 6), with generally lower numbers in crustaceans than in vertebrates and springtails. A Chi-squared test identified a statistically significant difference in the number of sites under selection between TM and MA+IM regions in traits EC, PC, and PM in mammals, as well as EC in *Pseudoniphargus* and vertebrates, PC in *Metacrangonyxs*, and PM in primates.

**Figure 6.**
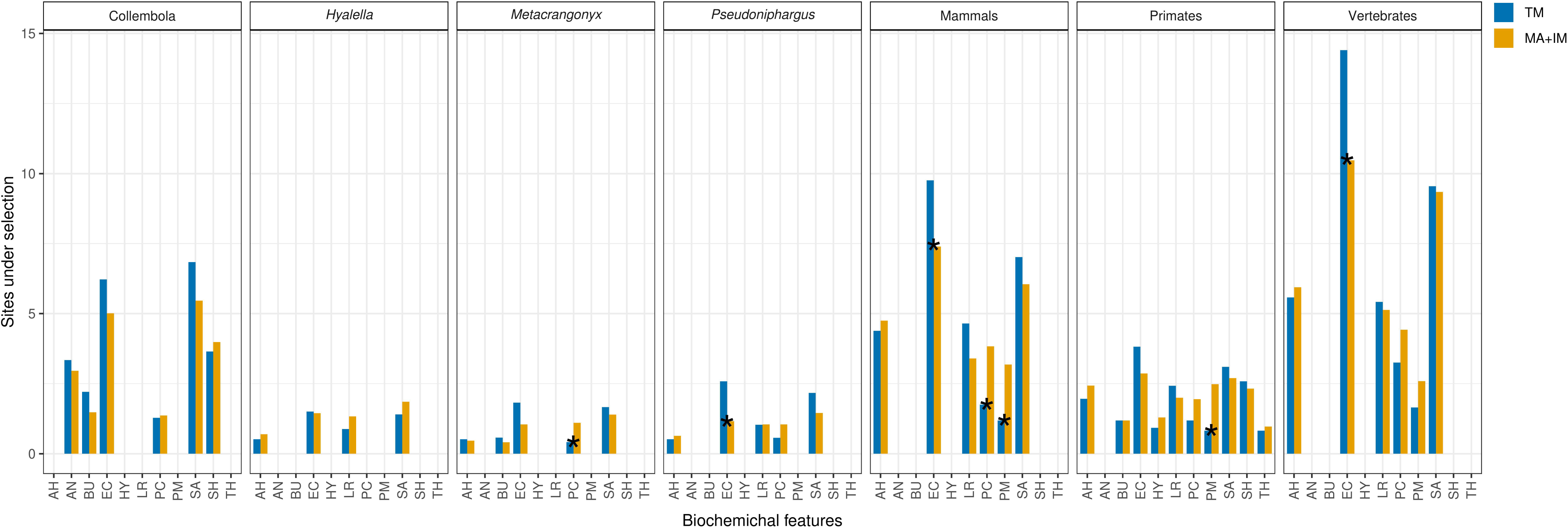
Number of amino-acid sites under selection for different biochemical features in TM vs. MA+IM domains. Asterisks indicate significant differences between TM and MA+IM regions (*p* ≤ 0.05). Biochemical features encoded as: Alpha-Helical tendencies (AH), Average Number of surrounding residues (AN), BUriedness (BU), Equilibrium constant ionization of COOH (EC), HydropathY (HY), Long-Range non-bonded energy (LR), Power to be at the C-terminal (PC), Power to be at the Middle of alpha-helix (PM), Solvent accessible reduction Ratio (SA), Surrounding Hydrophobicity (SH), and Thermodynamic transfer Hydrohphobicity (TH).

**Table 2.** Biochemical properties under positive selection for the seven datasets under scrutiny. . Symbol + indicates that the biochemical property is under selection.

|  | AH | AN | BU | EC | HY | LR | PC | PM | SA | SH | TH | total |
| --- | --- | --- | --- | --- | --- | --- | --- | --- | --- | --- | --- | --- |
| Collembola |  | + | + | + |  |  | + |  | + | + |  | 6 |
| *Hyalella* | + |  |  | + |  | + |  |  | + |  |  | 4 |
| *Metacrangonyx* | + |  | + | + |  |  | + |  | + |  |  | 5 |
| *Pseudoniphargus* | + |  |  | + |  | + | + |  | + |  |  | 5 |
| Mammals | + |  |  | + |  | + | + | + | + |  |  | 6 |
| Primates | + |  | + | + | + | + | + | + | + | + | + | 10 |
| Vertebrates | + |  |  | + |  | + | + | + | + |  |  | 6 |
| total | 6 | 1 | 3 | 7 | 1 | 5 | 6 | 3 | 7 | 2 | 1 |  |

Overall, a clear pattern does not emerge in terms of the specific evolutionary pressure at play in the different lineages, apart from a large prevalence of purifying over directional selection. Nevertheless, TM helices and MA+IM domains were shown to evolve under different selective pressures in all lineages, in the overall extent of positive and negative selection, as well as in the selection of specific biochemical processes.

## Discussion

Hitherto, the deployment of transmembrane region information in phylogenetic studies was limited to the assessment of orthology in protein families (Bjarnadóttir et al., 2006; Wojtkowska et al., 2012). In addition, structural information, alongside relative solvent accessibility and residue-residue contact information, was further deployed in bioinformatic tools specifically designed to improve the accuracy of multiple sequence alignments (e.g. Deng and Cheng, 2011). Here, we developed the new pipeline TRAMPO to deploy structural information (i.e. transmembrane helices, mitochondrial matrix and intermembrane domains) to improve partitioning in phylogenetic studies, enabling the creation of improved evolutionary models for tree inference.

While its core concept can be of diagonal interest, TRAMPO is specifically dedicated to the analysis of datasets consisting of nucleotide sequences of the 13 mitochondrial PCGs since they code for proteins with multiple transmembrane helices. While this may be seen as an outdated marker in the era of phylogenomics, we wish to stress that, with the democratization of sequencing/assembly/annotation tools for mitochondrial genomes (Dierckxsens et al., 2017; Bernt et al., 2013; Cucini et al., 2021), the use of complete mitochondrial genome datasets in phylogenetics is an active area of research, leading to many and important contributions especially in poorly studied taxa. As such, and because its orthology is easy to establish, any improvement in the dedicated analytical tools is likely to have a significant impact.

Our results confirmed the presence of well known base composition inequalities previously described across both codon positions (Pons et al., 2010; Cucini et al., 2020) and coding strands (Hassanin et al., 2005), inequalities that were, in turn, shown to have a significant impact in phylogenetic inference. Besides, we showed, for the first time, that these differences are also associated to specific secondary structure features. Specifically, we demonstrated that second codon sites of transmembrane helices (TM) display an extremely high T bias (Fig. 1, Fig. S1) relative to both matrix (MA) and intermembrane (IM) domains. Differences were also detected at the amino-acid level, with transmembrane helices harbouring a higher fraction of hydrophobic residues, and a lower fraction of acid, amide, and basic residues, than both matrix (MA) and intermembrane (IM) domains (Fig. 2). In addition, evidence was provided for differential selective pressures acting over TM helices with respect to MA and IM domains. Noteworthy, these patterns were observed in all the seven lineages studied here and therefore most likely represent a shared, diagonal feature of mitochondrial genomes across the animal life, and not an idiosyncrasy of a specific lineage. This observation significantly adds to the relevance of the results presented.

Partitioning of mitochondrial PCGs, generally by codon and by strand, during a phylogenetic analysis is common practice, and it was shown to produce significant improvements in phylogenetic reconstructions across many lineages of highly diverse arthropods, such as beetles (Timmermans et al., 2010), grasshoppers (Fenn et al., 2008), hymenopterans (Song et al., 2016), crustaceans (Bauzà-Ribot et al 2012), as well as in vertebrates, including birds (Powel et al., 2013) and bovids (Bover et al., 2019). Furthermore, these studies demonstrated that the optimization of partitioning schemes, following iterative merging based on a statistic criterion (AIC, AICc or BIC), leads to additional improvements in model accuracy, model fit, and the quality of the phylogenetic reconstruction as measured by global ML estimation (i.e. topology plus branch lengths).

Our results indicate that an additional level of partitioning, based on a third and novel criterion, i.e. outer-, inner- and trans-membrane regions, improved the fit of the evolutionary model in all seven lineages. This improvement, though to a lesser degree than partitioning according to codon and strand, indicates that the inclusion of secondary structure information in partitioning schemes is beneficial. Implementation of the PartitionFinder algorithm to further optimize that initial partitioning scheme, almost invariantly identified second codon sites of TM helices as independent charset blocks, confirming that those sites are indeed characterized by a distinct evolutionary pattern relative to other partitions (Fig. 3). The presence of a single gene encoded in the negative strand of vertebrate mitochondrial genomes precluded the formation of independent transmembrane partitions on this strand, even in the highly biased second codon sites of transmembrane helices. In fact, in vertebrates, sites on negative strand were merged into two partition only: first and second codon positions *vs.* third codon positions (Fig. 3). This outcome is nevertheless not totally unexpected given the nature of the heuristic greedy algorithm implemented in PartitionFinder that, in the absence of a sizeable contrasting signal, is designed to merge partitions. In the case of the vertebrate negative strand, with one single gene, the number of sites in each structural category is utterly small, and the associated phylogenetic signal accordingly limited, leading to the merging of sites irrespective of structural domains. The same does not, in fact, happen in the vertebrate positive strand, where 12 genes are encoded and where, accordingly, partitions include a larger number of sites and, arguably, hold a stronger phylogenetic signal.

Concerning the actual effect of secondary structure information on the resulting tree, it was shown by Panday and Braun (2020) that this can be substantial, and the phylogenetic signal embedded in secondary structures that may have a great impact especially on deep nodes. They applied a similar analytical approach, though at a different scale, to include secondary structure information in their phylogenenomic analysis, assigning sites to different secondary structure features (helices, sheets, and coils, buried or non-buried based on solvent accessibility). The phylogenetic analysis of exposed sites resulted in a tree with ctenophores as the sister group to all other animals, while buried sites suggested a sponge+ctenophore clade. Recoding of amino acids to reduce compositional biases, in turn, increased the agreement between exposed and buried sites, leading support to ctenophores as the sister group to all other animals.

In our analyses, the impact of partitioning based on secondary structures on the tree topology was noticeable in general, but to a different extent in the seven lineages. In two lineages, *Hyalella* and *Pseudoniphargus*, and despite sizeable differences in likelihood values, different partition schemes did not result in topological changes. In springtails, mammals and primates, trees clustered into two tree islands, with low internal distances. Finally, the analysis of two lineages resulted in generally large inter-tree distances among trees obtained using different partition schemes (Fig. 4). Many clusters were observed in the *Metacrangonyx* tree space, and each partition resulted in a different tree for vertebrates (Fig. 4). Interestingly, the analysis of the most recent lineages (*Hyalella*, 25 Ma, Zapelloni et al., 2021; *Pseudoniphargus,* 53 Ma, Stokkan et al., 2018; Primates, 74 Ma, Pozzi et al., 2014) resulted in one single, or a tightly-knit set of trees, with low distances across. Analyses of older datasets, in turn, resulted in a wider tree space, with different partition schemes that support different, and sometimes fairly distant, trees (Fig. 4).

Notably, this does not seem to be connected with the number of species in each dataset, as it is similarly observed in *Metacrangonyx* (23 species) and vertebrates (150). Furthermore, and despite sizeable tree distances, topologies were still congruent in all seven lineages (negative AU-test) suggesting that these differences mostly arise due to suboptimal signal and not due to strong contrasting signals among partitioning schemes.

Our interpretation relates to the phylogenetic signal that characterize the three codon positions over different time frames. Partitioning by domain appears to be especially relevant for second codon positions, that are characterized by a different base composition in transmembrane helices compared to matrix and intermembrane domains (Figs. 1-3). As such, the effects of partitioning by domain are most likely driven by second codon positions. These latter positions are characterized by a lower substitution rate compared to first, and especially third, codon positions. Wherefore, third and first codon positions dominate over shorter phylogenetic times, whereas second codon positions are expected to become the most influential over longer time frames. Consequently, it can be hypothesized that the observed inequality among second codon positions in TM vs. MA+IM domains becomes evident, and take its effects, over longer time frames. This effect may be further exacerbated in *Metacrangonyx* that was shown to evolve under an accelerated rate of evolution (Bauzà-Ribot et al., 2012; Pons et al., 2019).

As expected, most mitochondrial protein sites were under purifying selection and few positions were identified as under positive selection, without a clear pattern across lineages. Sites under negative selection were 10- to 100-fold more abundant that sites under positive selection. Concerning positive selection, the number of amino acid sites under episodic diversifying positive selection was about 10-fold higher than sites under pervasive positive diversifying selection. Crustaceans lineages appear to have a lower number of sites under positive selection relative to springtails and the three vertebrate lineages (Fig. 5). It cannot nevertheless be excluded that this was the outcome of an ascertainment bias associated to the lower number of species in crustacean datasets that, most likely, make the analysis less effective. Concerning differences among structural domains, an excess of sites under episodic diversifying positive selection was observed in mammals and vertebrates in transmembrane regions, whereas no significant difference is observed in pervasive positive diversifying selection. Marginal but statistically significant differences were also observed in the number of sites under negative selection in three lineages: *Hyalella* and *Metacrangonyx* (where negative selection significantly influences MA+IM domains) and vertebrates (where negative selection especially affects TM helices; Fig. 5c).

Moving from the general assessment of positive selection to the assessment of selection over specific biochemical properties (Table 2), some additional trends can be extrapolated. Some biochemical properties appear to be under positive selection in all, or almost all, groups under study (AH: alpha-helical tendency; EC: equilibrium constant ionization of COOH; PC: power to be at the C-terminal; SA: solvent accessible reduction ratio). This lends credibility to this observation and suggests these as general trends of mitochondrial proteins not associated to a specific lineage. To be singled out is also PM (power to be at the middle of an alpha-helix) that is identified as being under positive selection exclusively in vertebrate taxa, potentially indicating a trend along the vertebrate branch of the animal tree. Regarding differences among structural domains (TM vs. MA+IM) in terms of specific biochemical properties under selection, seven instances were identified that are characterized by significant inequalities. These are concentrated in three biochemical properties: equilibrium constant ionization of COOH (EC), with an excess in TM regions, as well as power to be at the middle of an alpha-helix (PM) and power to be at the C-terminal (PC), with a deficiency in TM regions. Notably, five of these outcomes are concentrated in the three vertebrate lineages that were previously singled out for a significantly higher number of sites under selection, overall, in TM vs. MA+IM domains (Fig. 6). In summary, while not pervasive, nor structured with a clearly recognizable pattern by lineage or by biochemical property, there is indication of ongoing selection in the seven datasets (Fig. 6). Most importantly, selection is shown to affect transmembrane domains differently compared to matrix and intermembrane domains. This is visible in terms of the large number of sites in TM vs. MA+IM under positive selection and a different nature of the biochemical properties that are over/under-represented in TM vs. MA+IM.

In other studies some biochemical features under positive selection were related to metabolic adaptation and potential effects of historical climate events and local adaptation. For instance, the equilibrium constant (ionization of COOH) is under selection in the mitochondrial cytochrome c oxidase complex in pikas, aiding adaptation to high elevations and influencing COX complex efficiency in lowland species for better heat production (Solari and Hadly, 2018). Similar positive selection for this property, along with hydropathy and molecular volume, was found in turbellarians (Li et al., 2024), suggesting a crucial role in adapting to low oxygen environments, while intertidal chitons (Dhar et al., 2021) and the killifish *Aphanius fasciatus* (Pappalardo et al., 2024) were related to eurythermal and euryhaline adaptations. Here we found that some lineages, particularly those with a larger number of species, showed more amino acid sites under strong positive selection for certain biochemical properties. Notably, properties like Power to be at the C-terminal (PC), Equilibrium constant ionization of COOH (EC), and Power to be at the Middle of alpha-helix (PM) exhibited differences between TM and MA+IM regions, with some results possibly influenced by sample size and test sensitivity. Noteworthy, the only feature found under positive selection in the three vertebrate lineages PM (3) also found difference between TM and MA+IM. In *Metacrangonyx* and *Pseudoniphargus*, these biochemical features under selection could be related to adaptation to the harsh stygobiont environment characterized by oligotrophy, complete darkness, and low oxygen content. On the other hand, *Hyalella* species that inhabit in lakes does not show distinct selective pressure between TM helices and MA+IM domains. Nonetheless, this hypothesis requires further investigation that is beyond the scope of this study.

Based on the observations conducted here, we recommend the partitioning of sites of mitochondrial PCGs according to structural domain (trans-membrane region, matrix and intermembrane domains), alongside by codon and by strand, as a prerequisite for phylogenetic analysis. This appears to be particularly important in old lineages (> 50 Ma). Partitioning according to structural domains can be efficiently accomplished using the novel TRAMPO script, available from a dedicated GitHub page https://github.com/dbajpp0/TRAMPO.

## Acknowledgments

Joan Pons is indebted to Professor Carlos Juan for the long discussions we had, more than ten years ago, about the idea of using secondary structures of mitochondrial proteins in a phylogenetic framework. Trempó (pronounced trəm[po in Catalan), incorrectly written trampó in Spanish, is a simple summer salad typical from Mallorca made with tomatoes, pale green peppers and onions, dressed with salt and extra virgin olive oil, that blends the colours of the Italian flag.

## Supplementary Material

**Figure S1.** Scatter plots of Askew (x-axis) vs G-skew (y-axis) for different domains (IM, MA, TM), strand (positive and negative) and codon position for all datasets: Collembola, *Hyalella*, *Metacrangonyx*, *Pseudoniphargus*, Mammals, Primates, and Vertebrates, page 1 to 7, respectively. Partitions are named as ‘strand_domain_codon position’. Due to the presence of only one gene encoded in the negative chain of vertebrates, the three vertebrate panels don’t display strand differences.

**Figure S2.** Box plots of G+C frequency for different domains (IM, MA, TM), strand (positive and negative) and codon position for all datasets: Collembola, *Hyalella*, *Metacrangonyx*, *Pseudoniphargus*, Mammals, Primates, and Vertebrates, page 1 to 7, respectively. Partitions are named as ‘strand_domain’ and the tree codon positions are depicted as different panels. Due to the presence of only one gene encoded in the negative chain of vertebrates, the three vertebrate panels don’t display strand differences.

**Figure S3.** Frequency of amino-acids in different chemical classes in different domains (IM, MA, TM) and strands (positive and negative): Collembola, *Hyalella*, *Metacrangonyx*, *Pseudoniphargus*, Mammals, Primates, and Vertebrates, page 1 to 7, respectively. Partitions are named as ‘domain_stand’. Due to the presence of only one gene encoded in the negative chain of vertebrates, the tree vertebrate panels don’t display strand differences.

**Table S1.** AICc values associated with different partition schemes following merging/optimized with Partition Finder. AICc values are given for the first row only while the difference in percentage, relative to the first one, is given for the following rows. Colours indicate those partitions schemes that retrieved second codon sites of TM helices as independent partitions after optimization in Partition Finder: from positive strand only (orange), each strand as single independent sets (blue), and both strands merged in one (red).

**Table S2.** Charsets resulting from PartitionFinder merging and optimization of initial character sets with 18 12, and 9 partitions.

**Data_S1.** Concatenated alignment of thee 13 mitochondrial protein coding genes as well as seven different partitions schemes built with different combinations of functional sets (by codon, by strand and by secondary structure) for the seven lineages studied here. This file also includes trees, AICc values and PartitionFinder results obtained in IQ-TREE2.

