## Supplementary figures and images for "Integrating Secondary Structure Information Enhances Phylogenetic Signal in Mitochondrial Protein Coding Genes"

### Supplemetal FigureS1

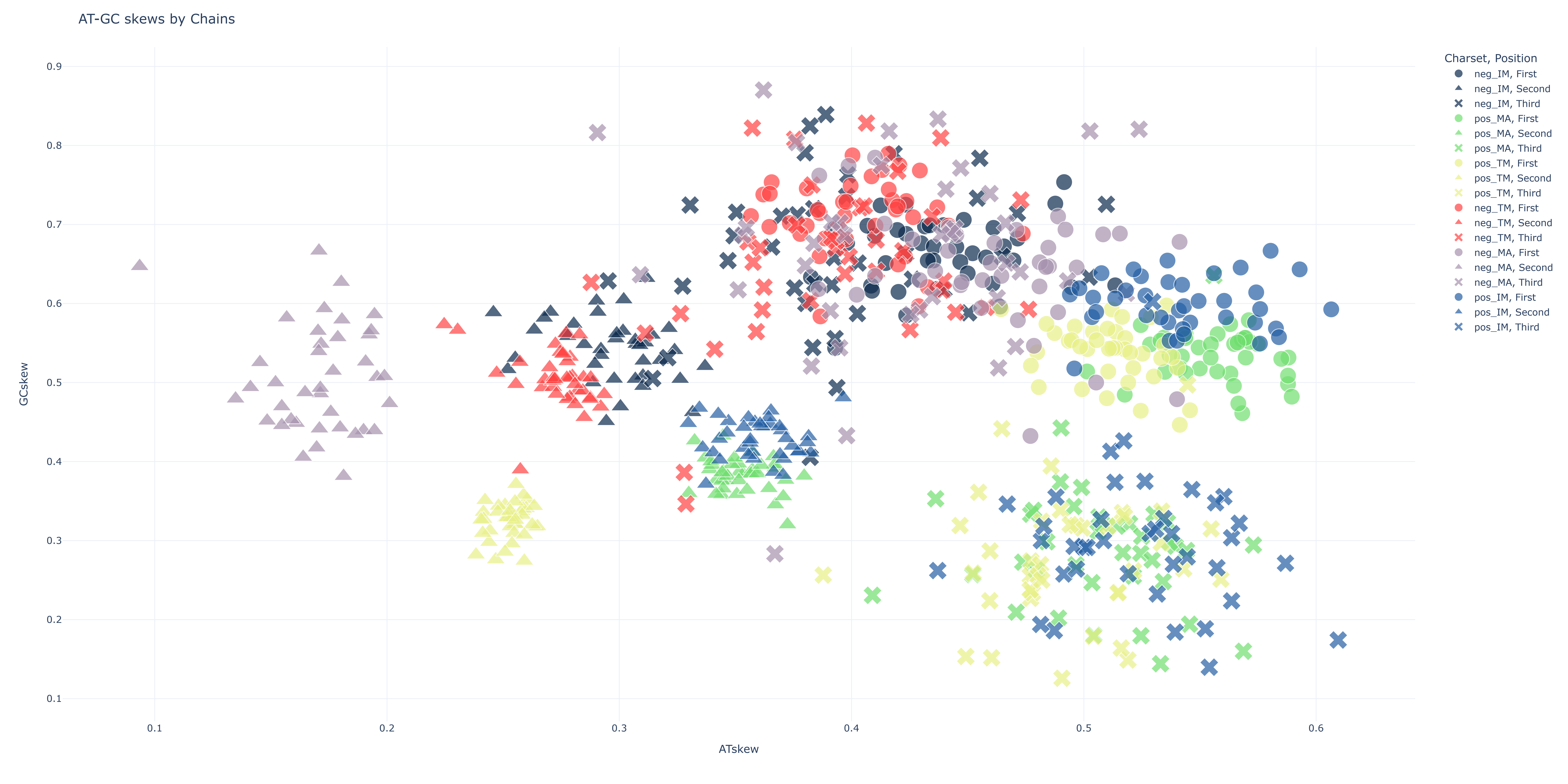

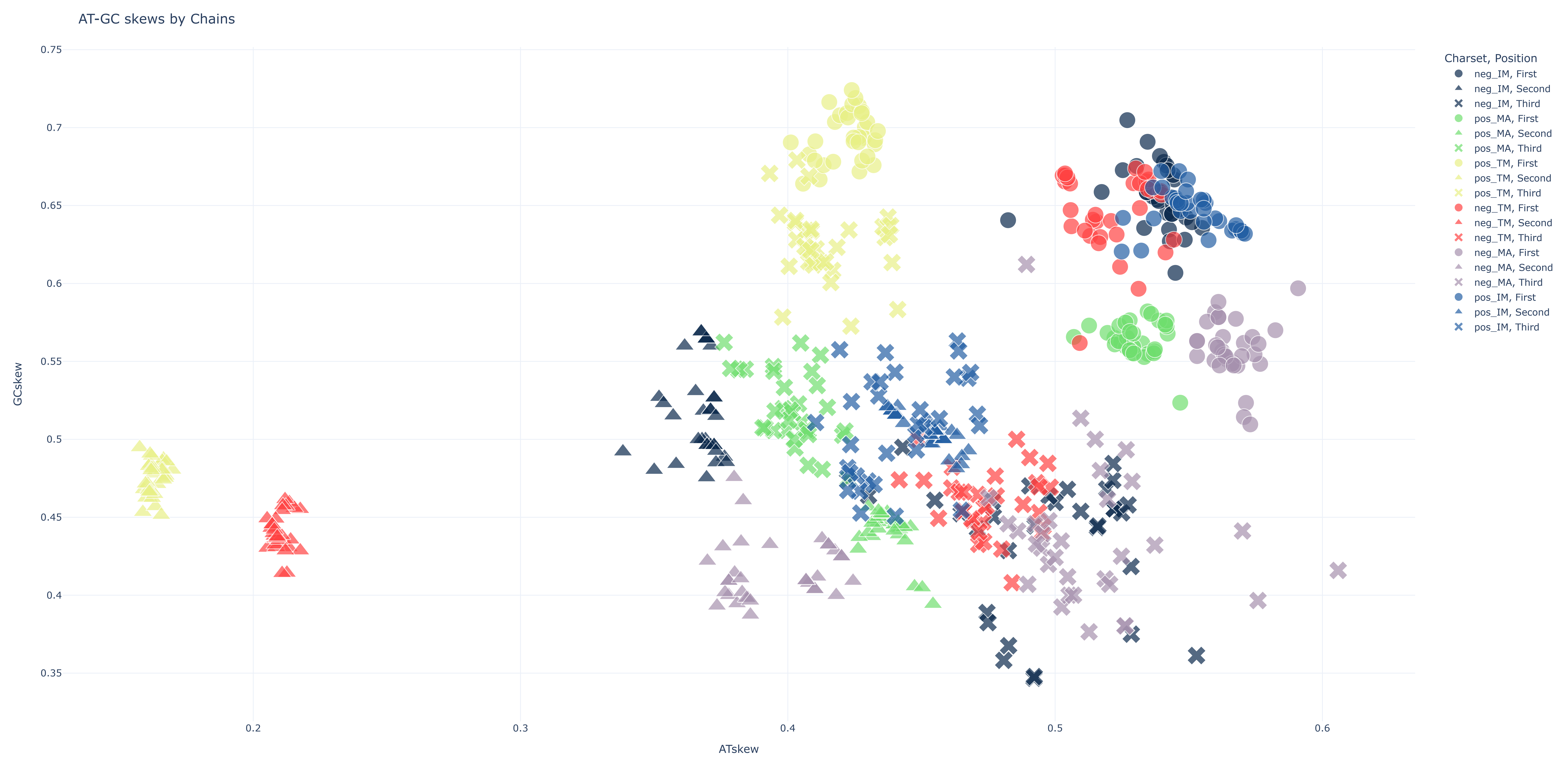

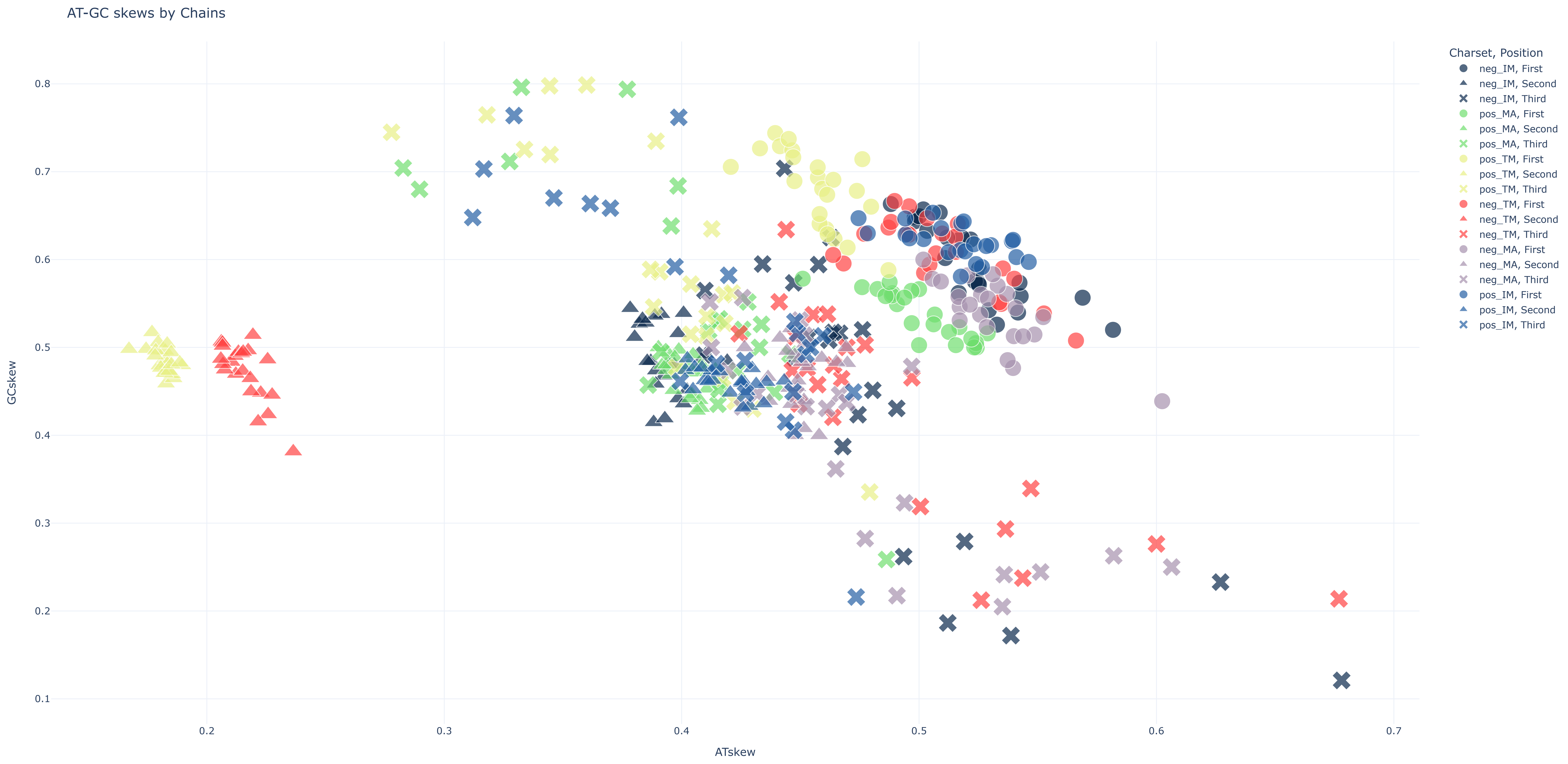

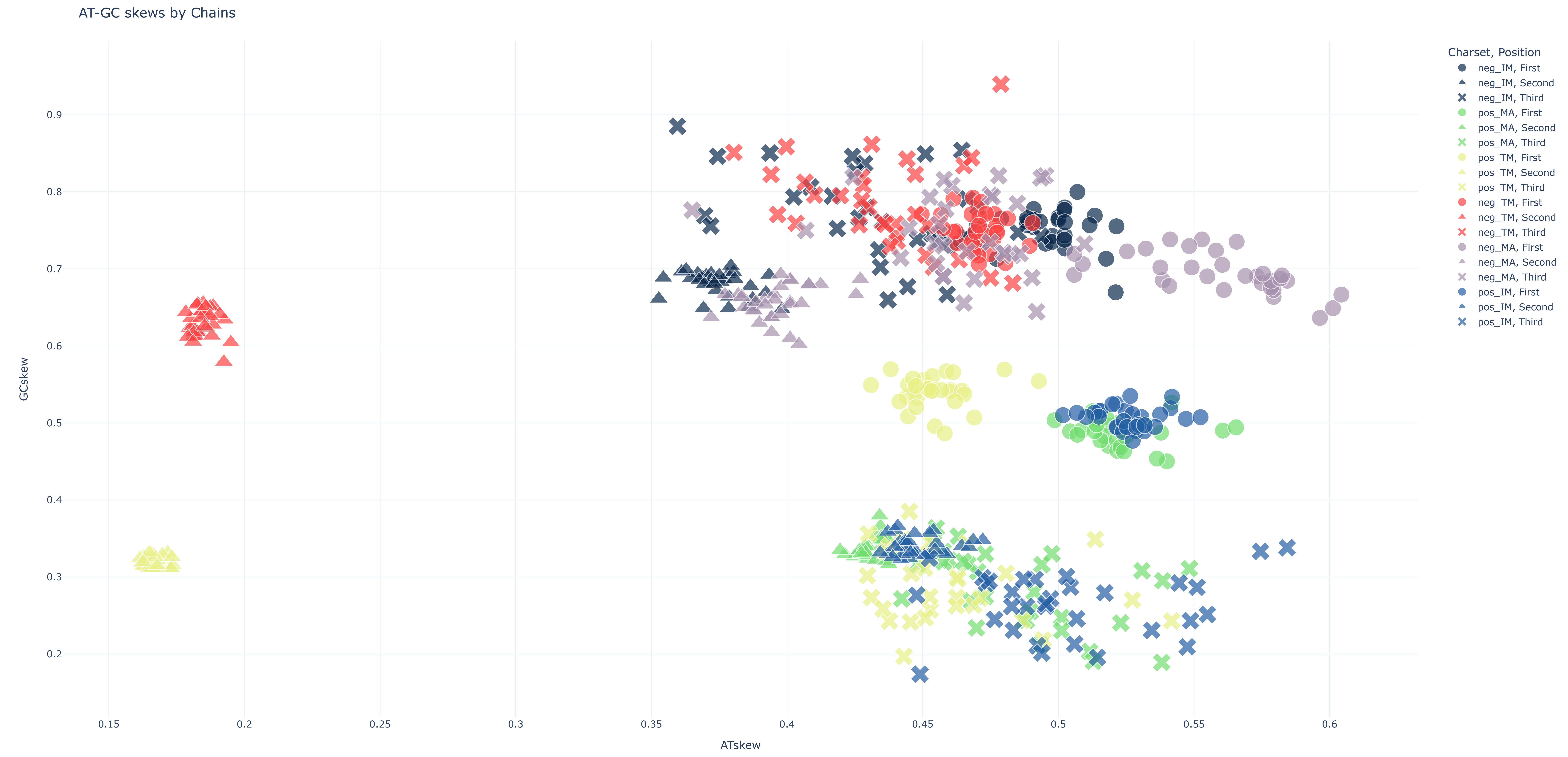

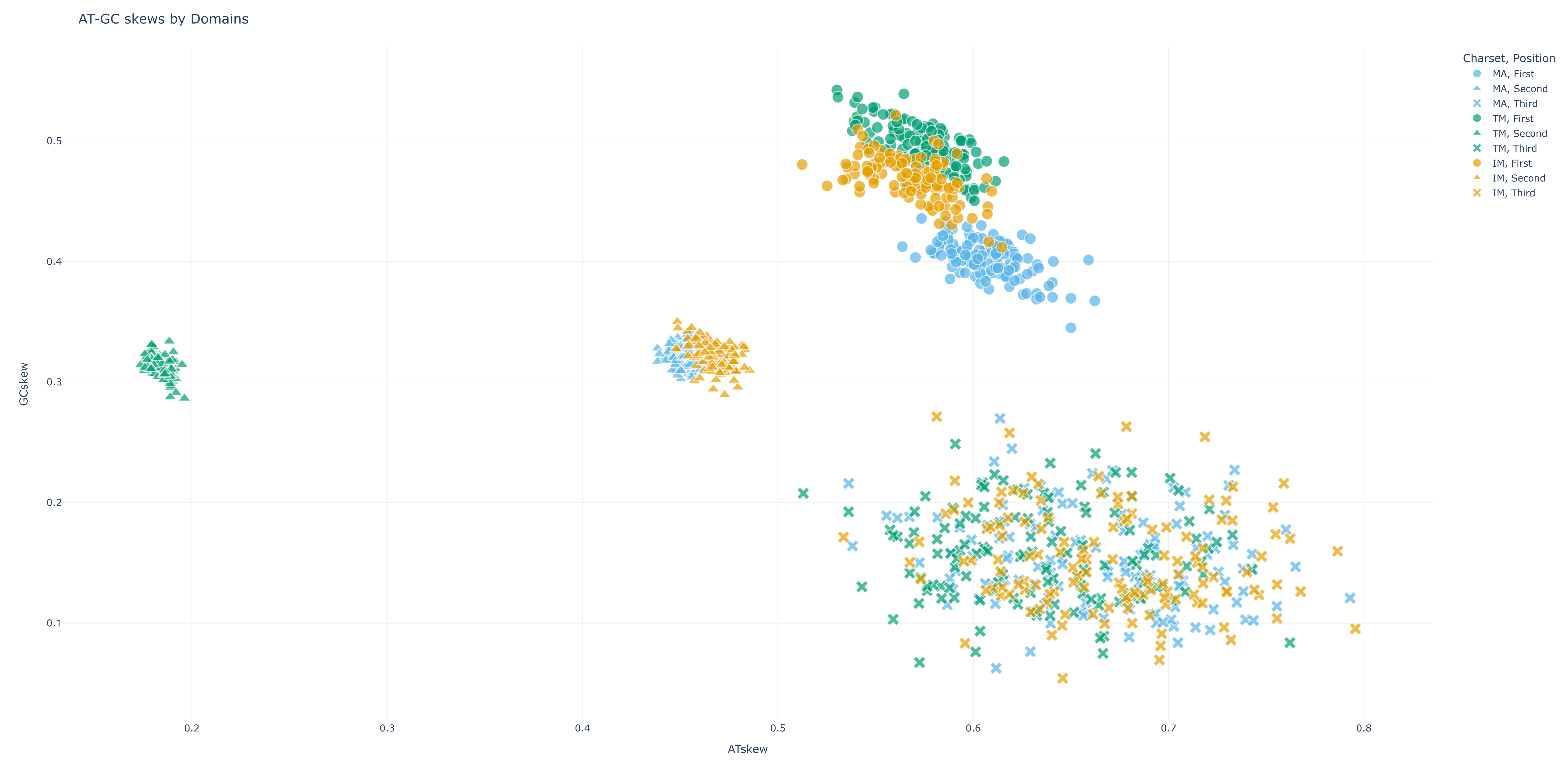

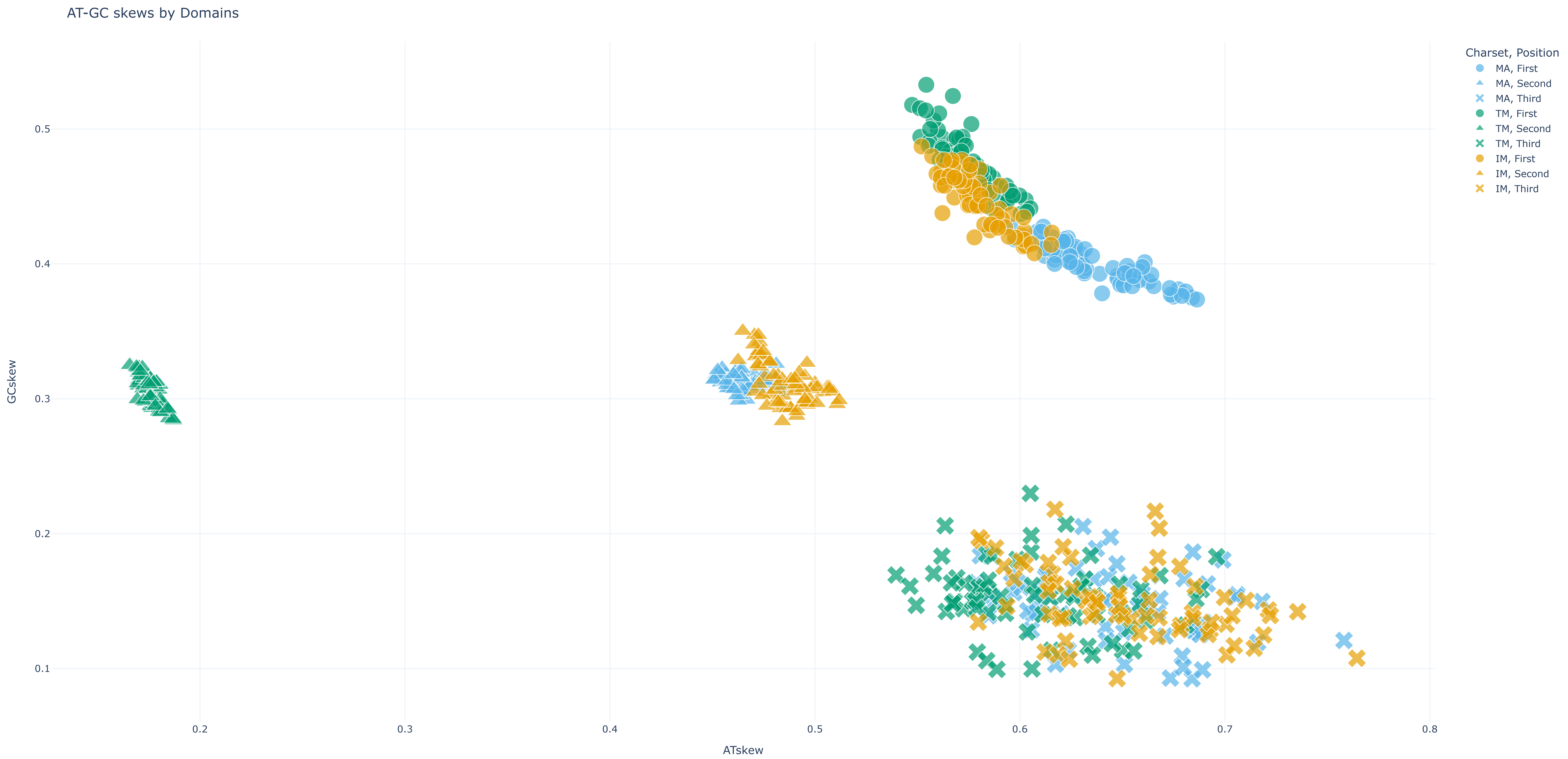


Loading [MathJax]/extensions/MathMenu.js

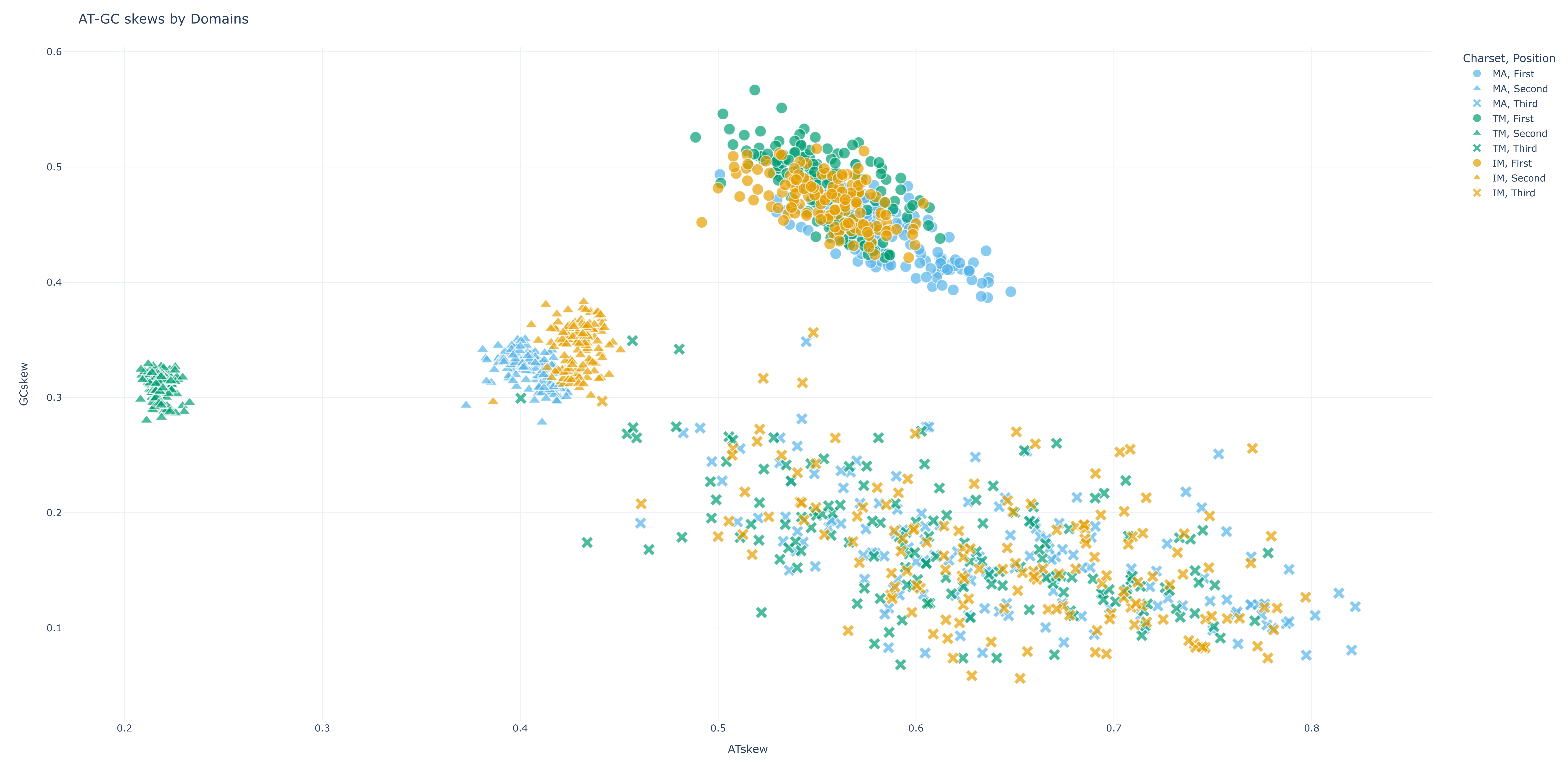
