## Supplementary material for "Integrating Secondary Structure Information Enhances Phylogenetic Signal in Mitochondrial Protein Coding Genes": Supplemetal FigureS2

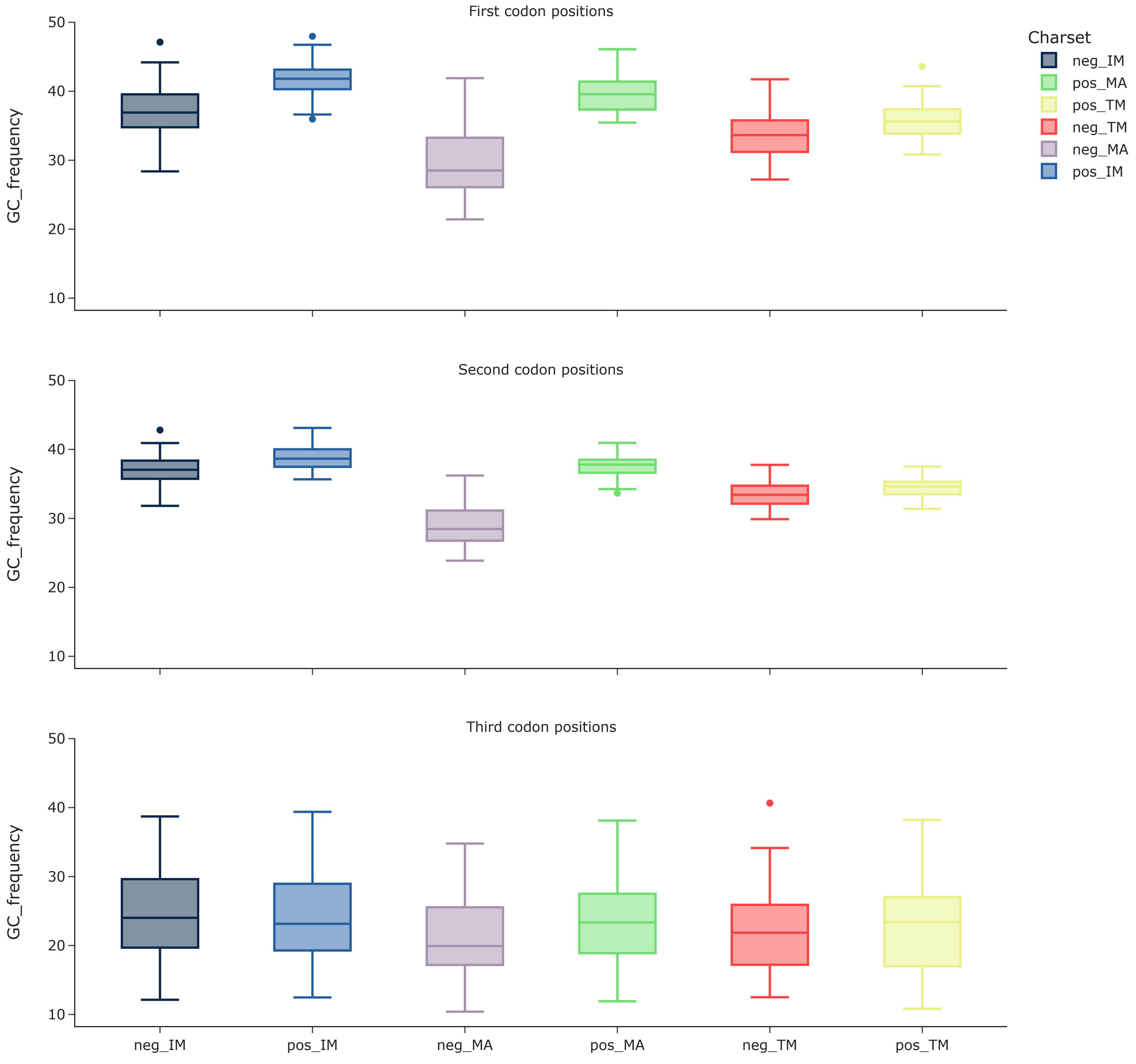


Loading [MathJax]/extensions/MathMenu.js

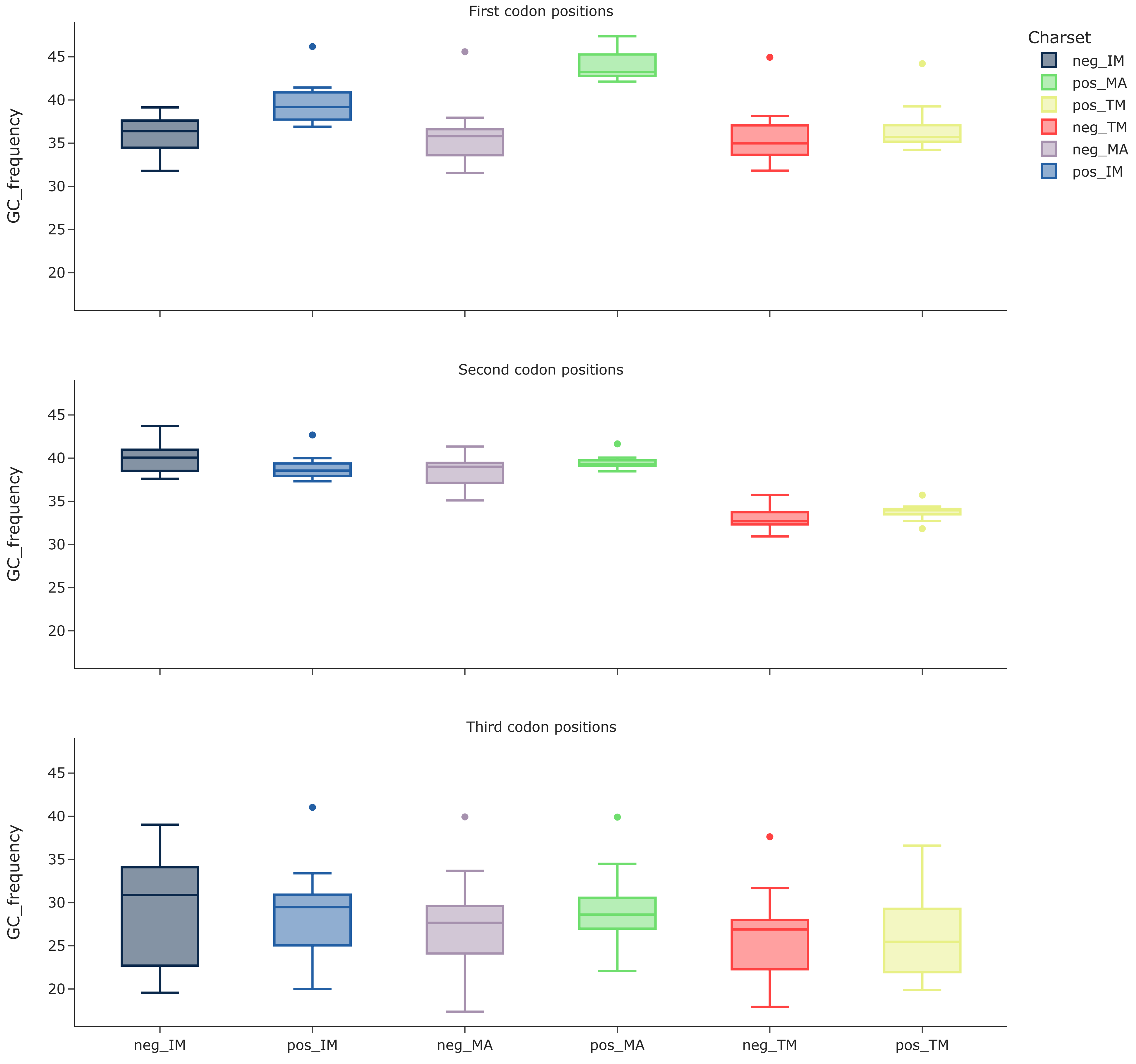


Loading [MathJax]/extensions/MathMenu.js

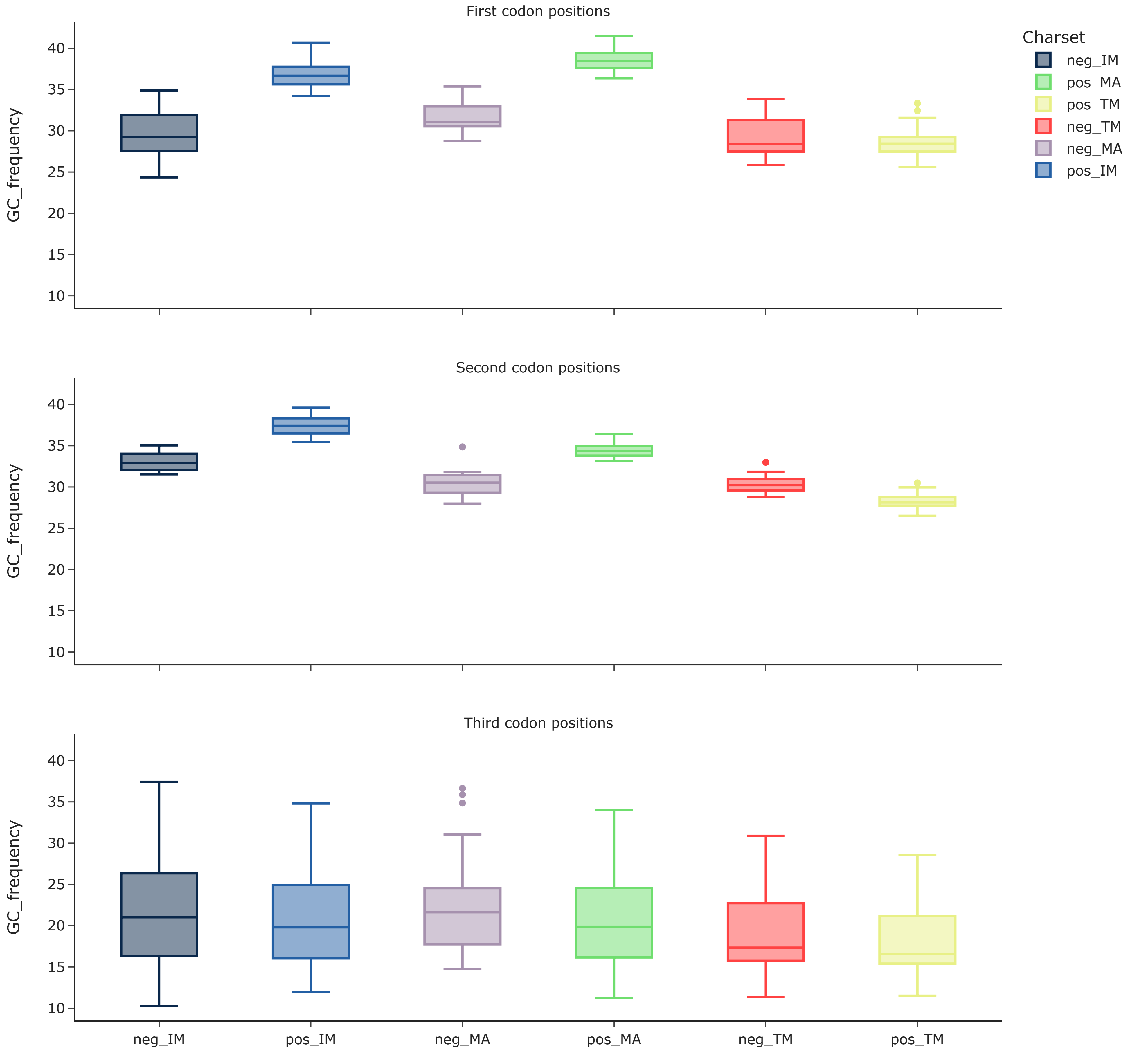


Loading [MathJax]/extensions/MathMenu.js

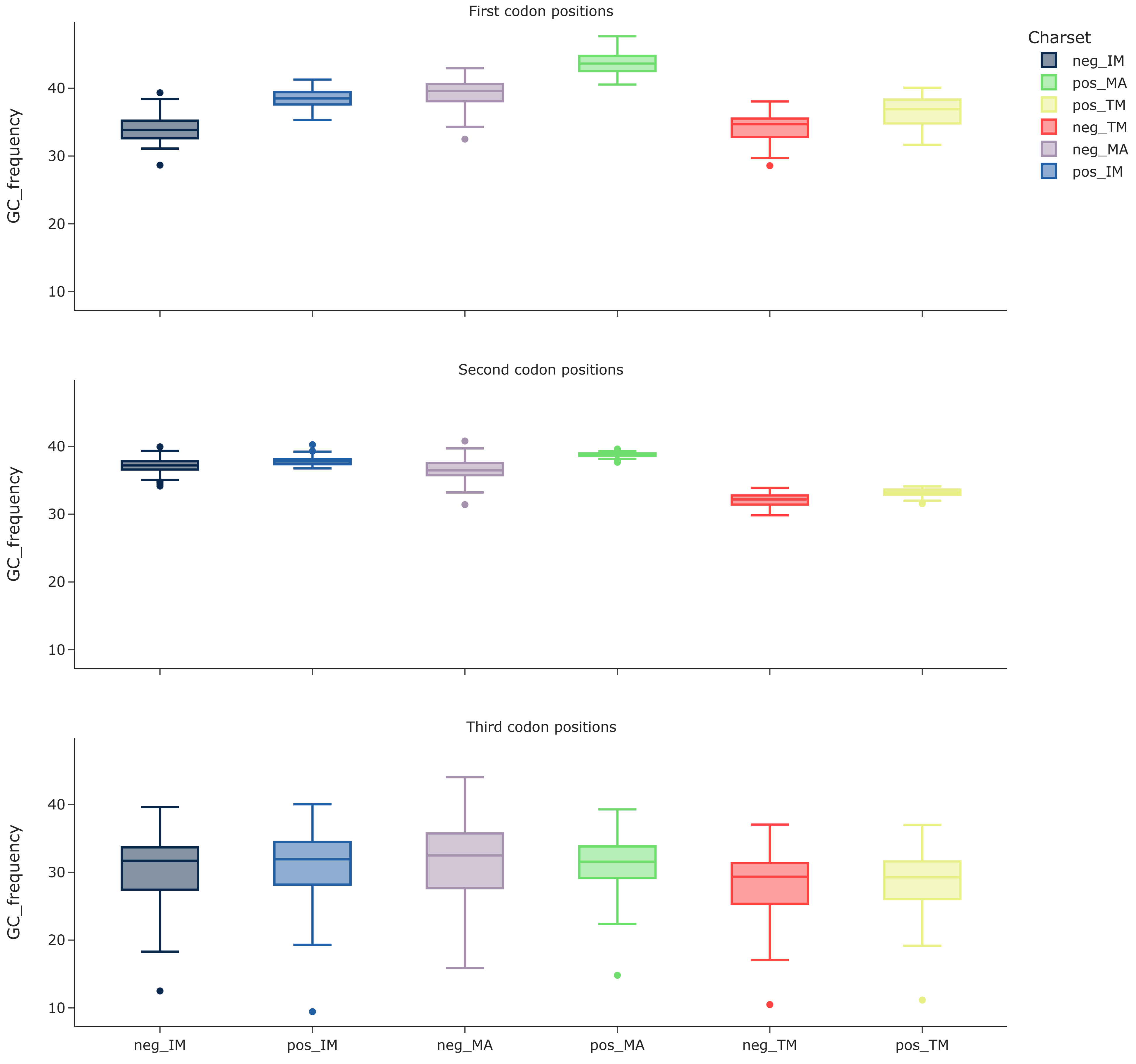


Loading [MathJax]/extensions/MathMenu.js

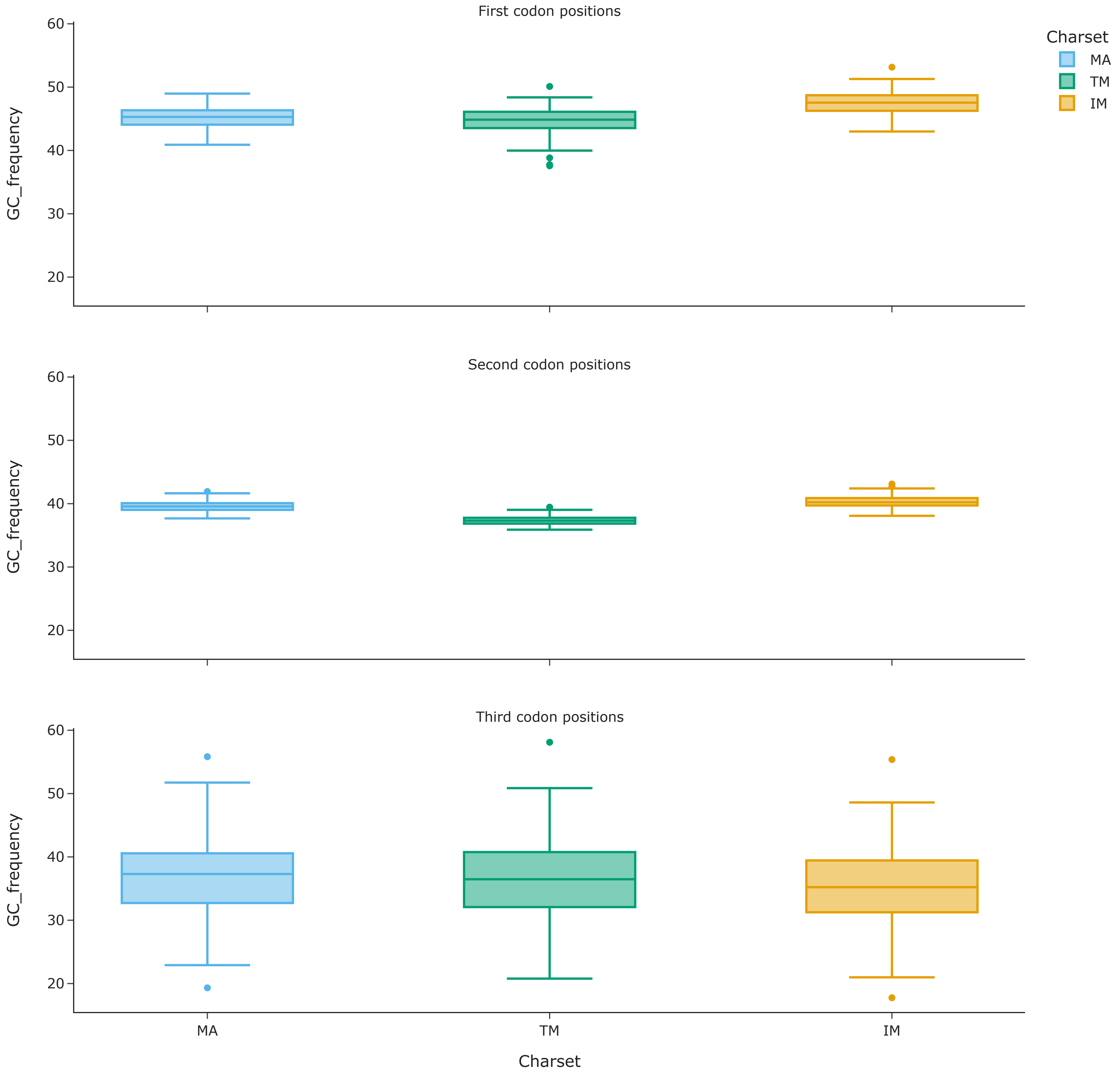


Loading [MathJax]/extensions/MathMenu.js

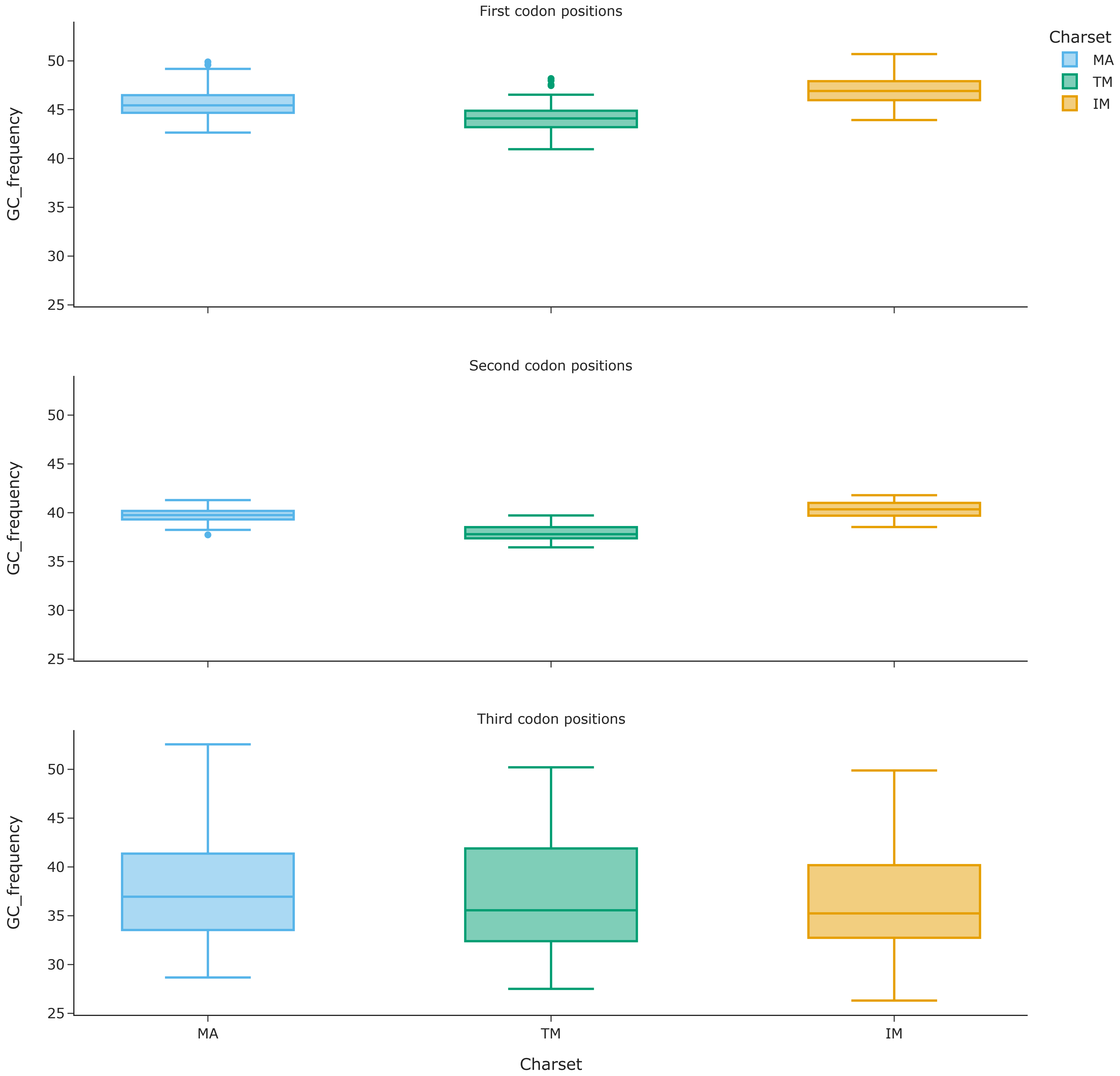


Loading [MathJax]/extensions/MathMenu.js

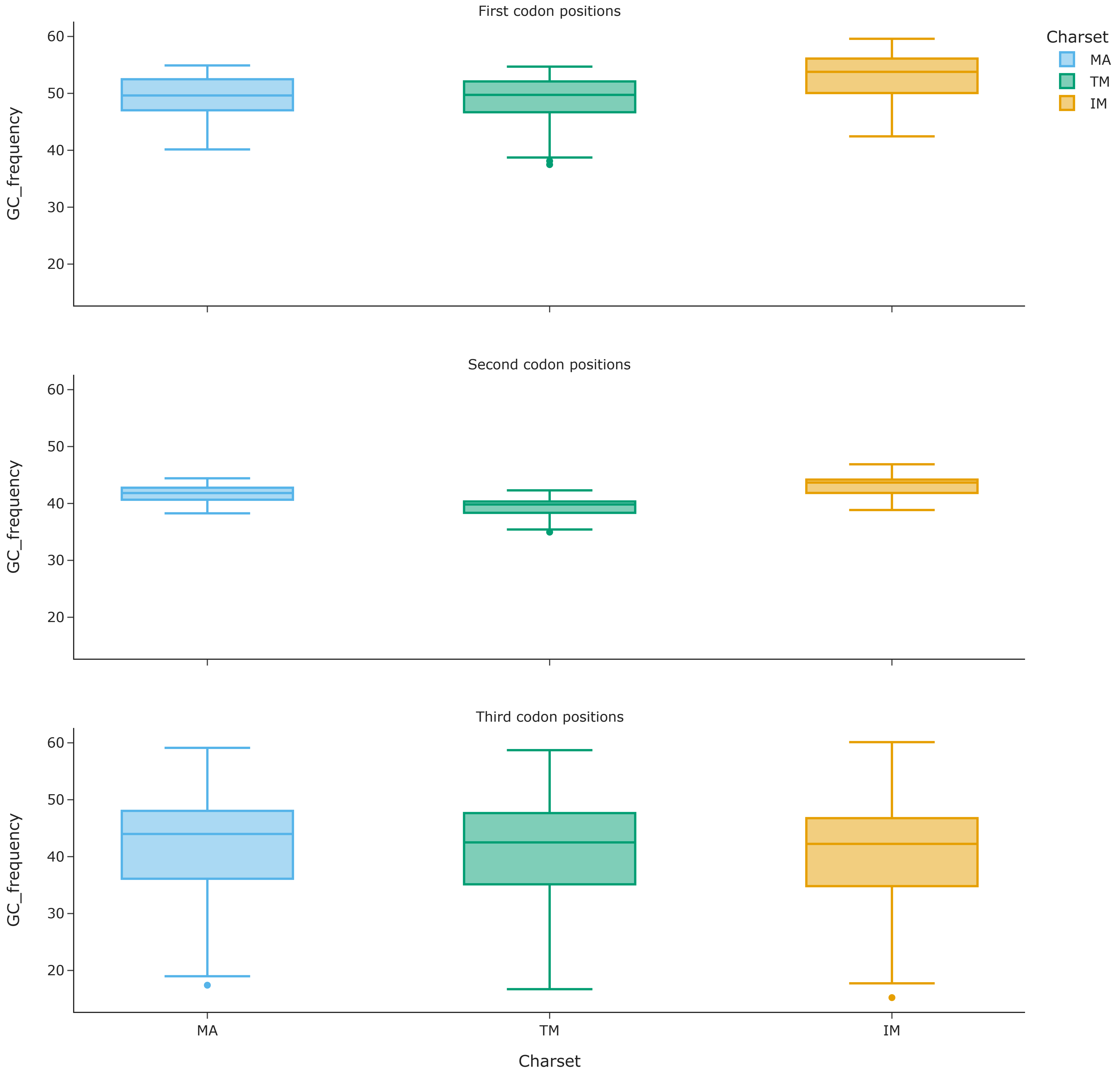


Loading [MathJax]/extensions/MathMenu.js
