## Supplementary material for "Integrating Secondary Structure Information Enhances Phylogenetic Signal in Mitochondrial Protein Coding Genes": Supplemetal FigureS3

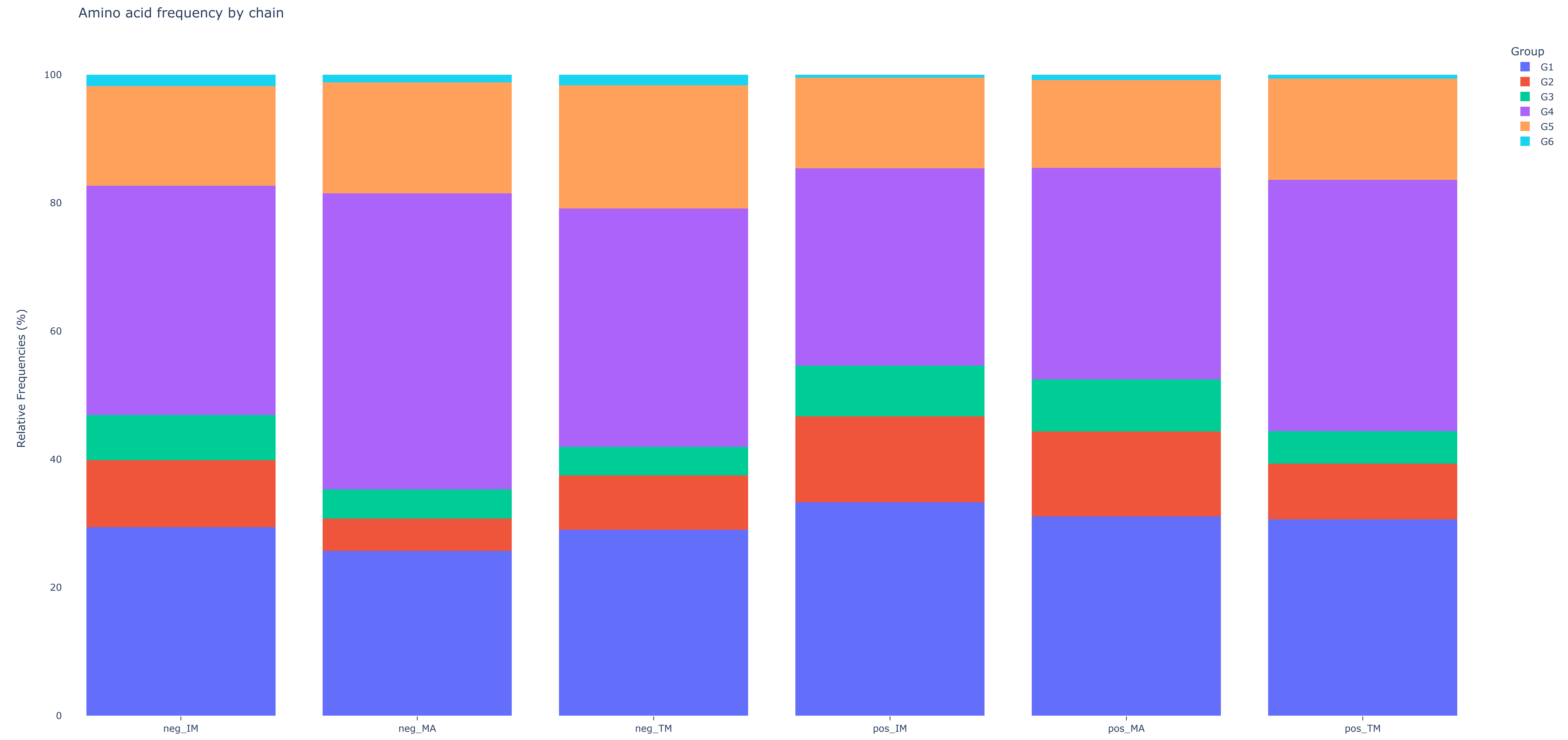

Loading [MathJax]/extensions/MathMenu.js

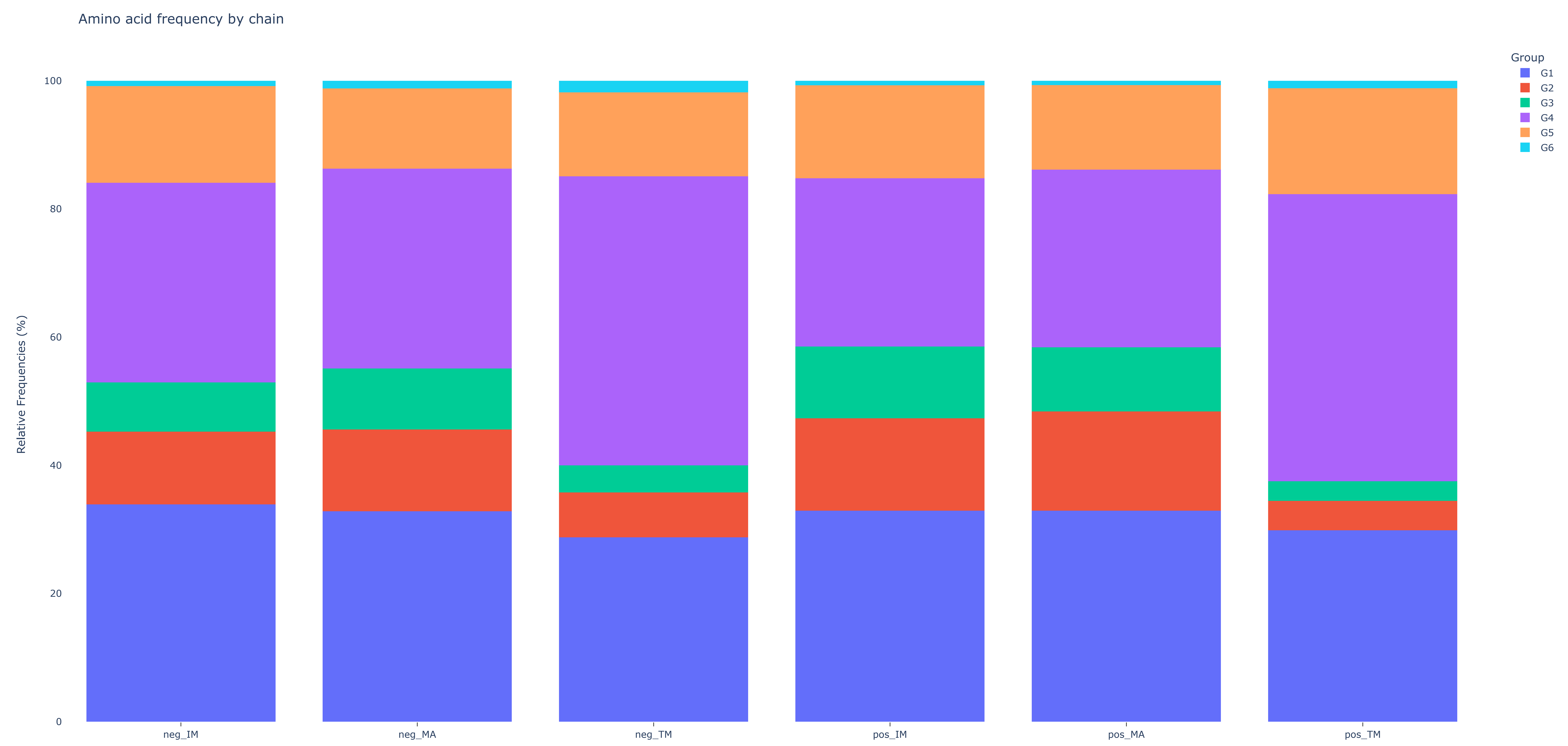


Loading [MathJax]/extensions/MathMenu.js

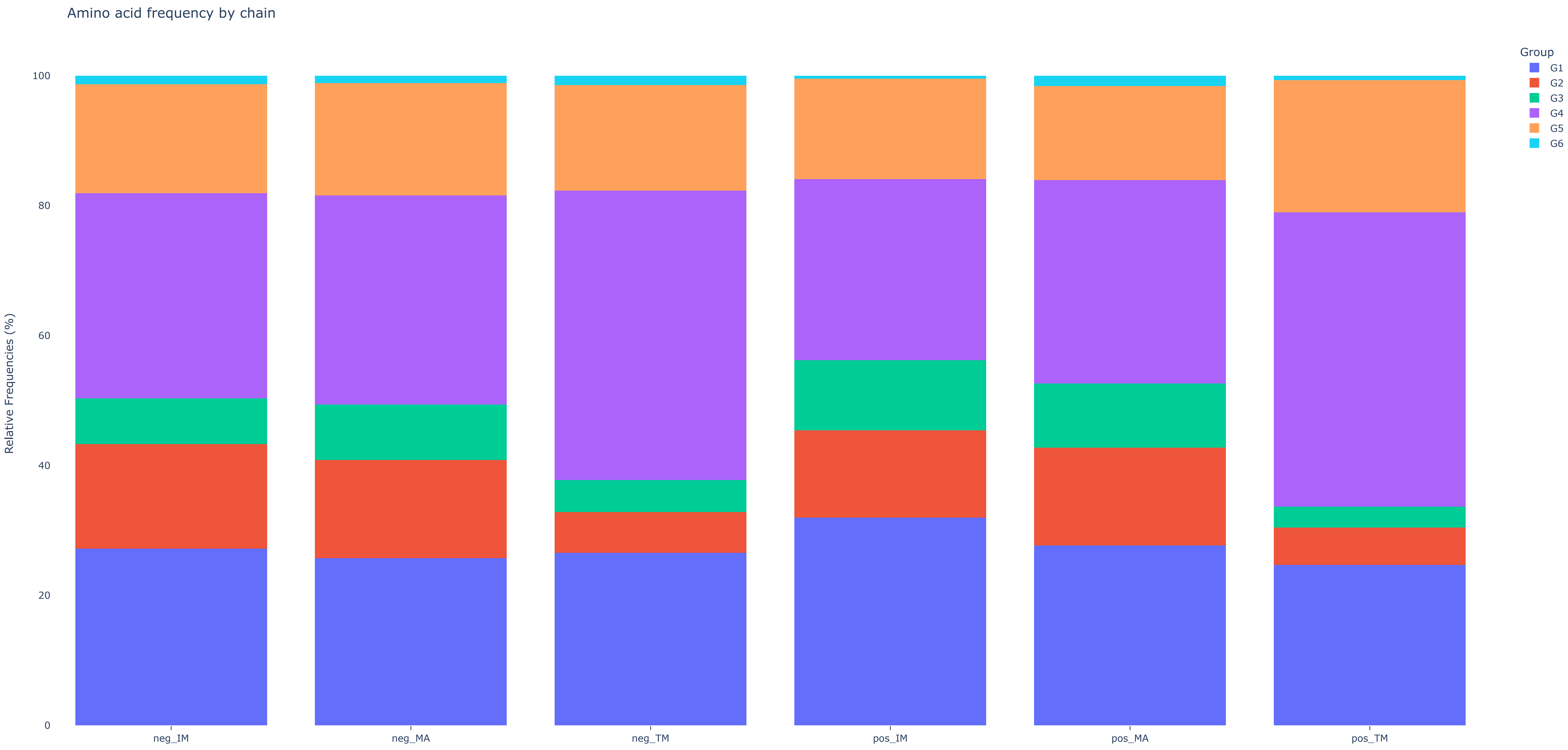

Loading [MathJax]/extensions/MathMenu.js

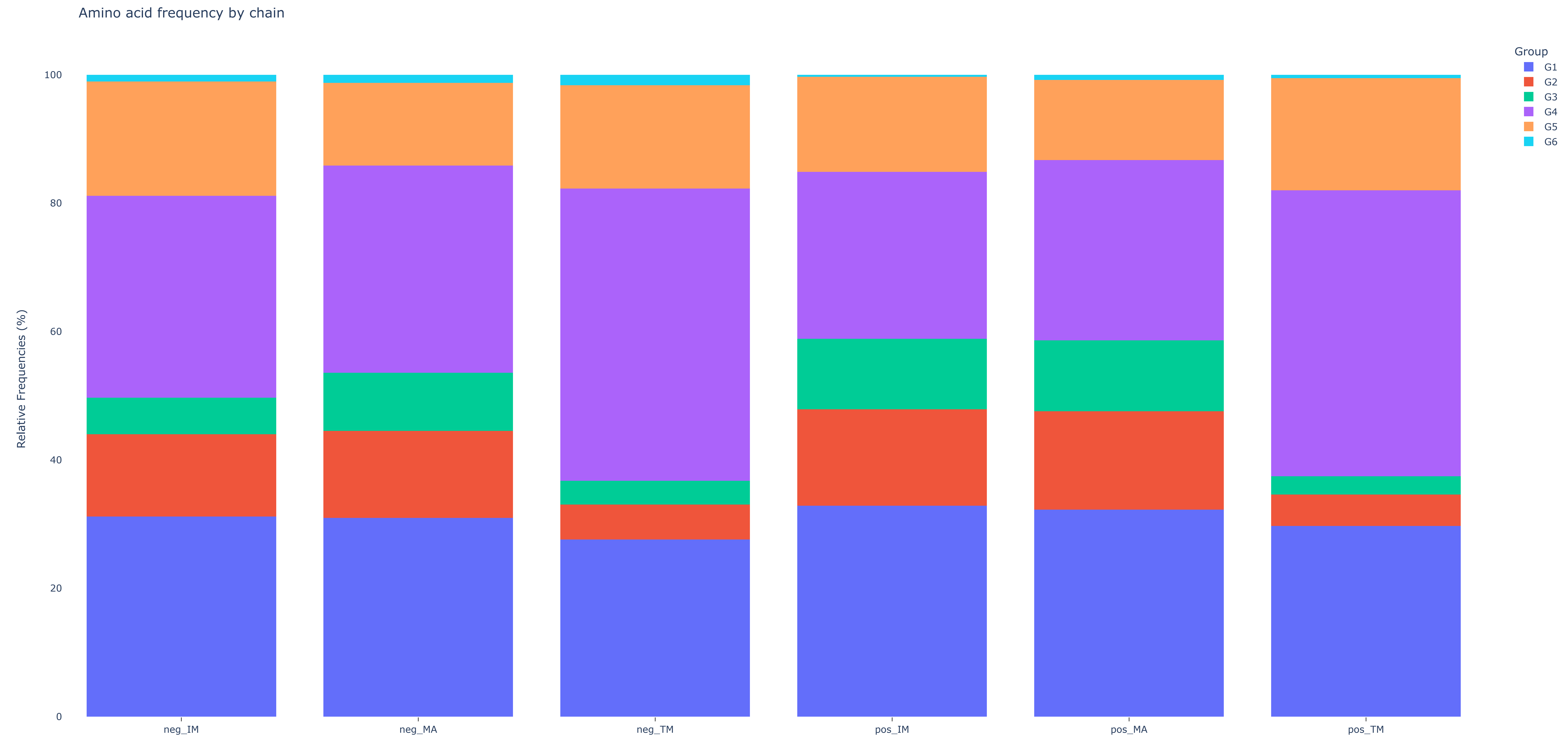


Loading [MathJax]/extensions/MathMenu.js

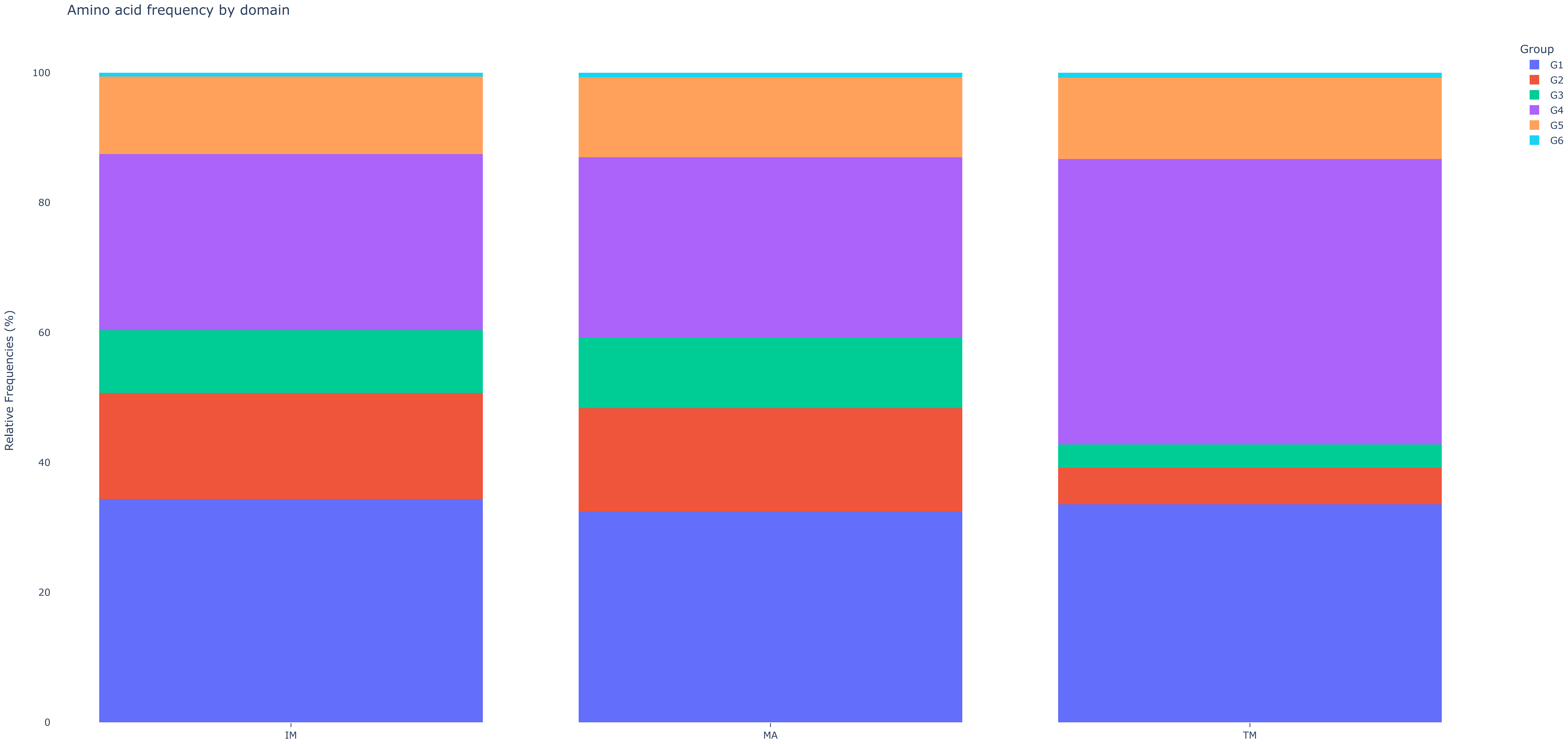

Loading [MathJax]/extensions/MathMenu.js

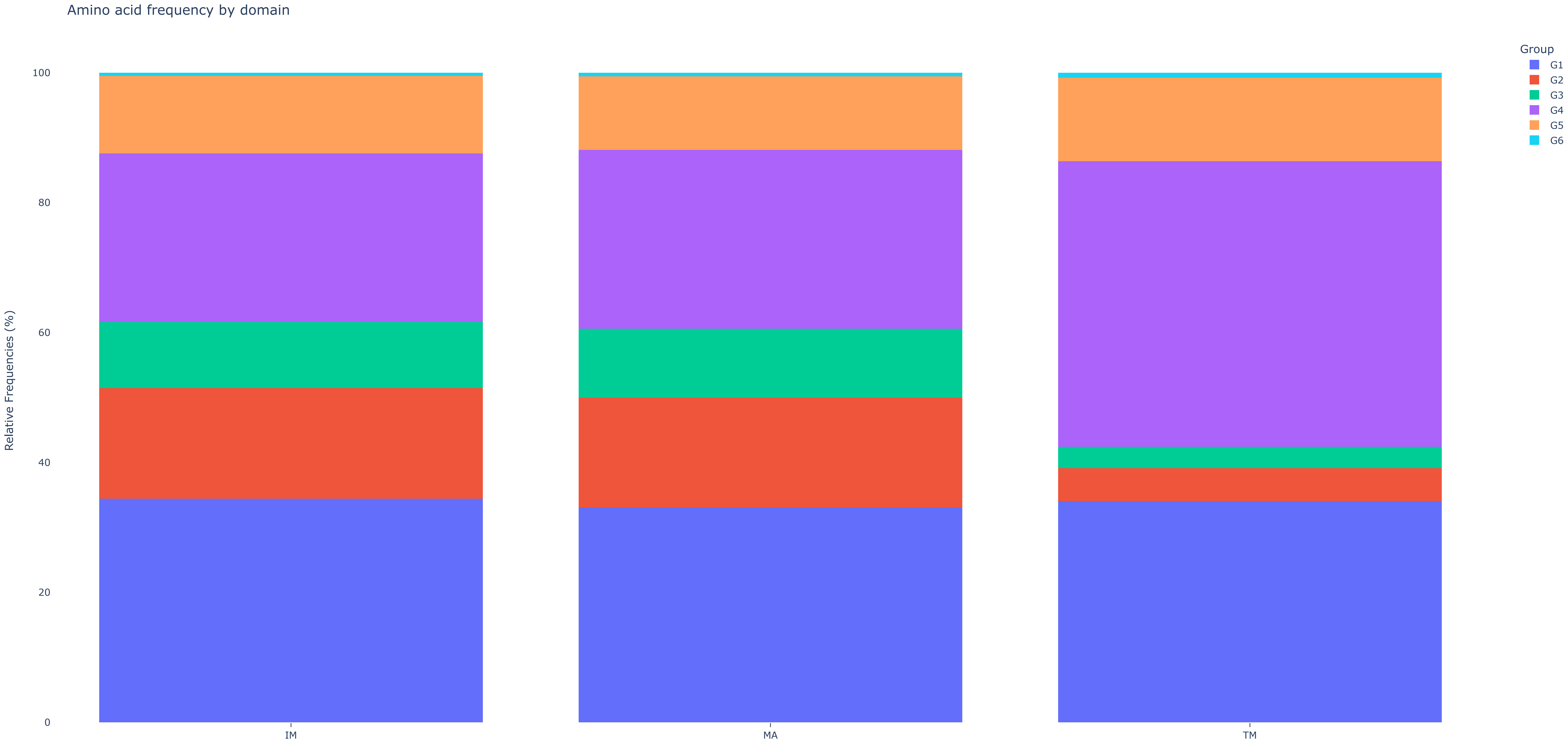


Loading [MathJax]/extensions/MathMenu.js

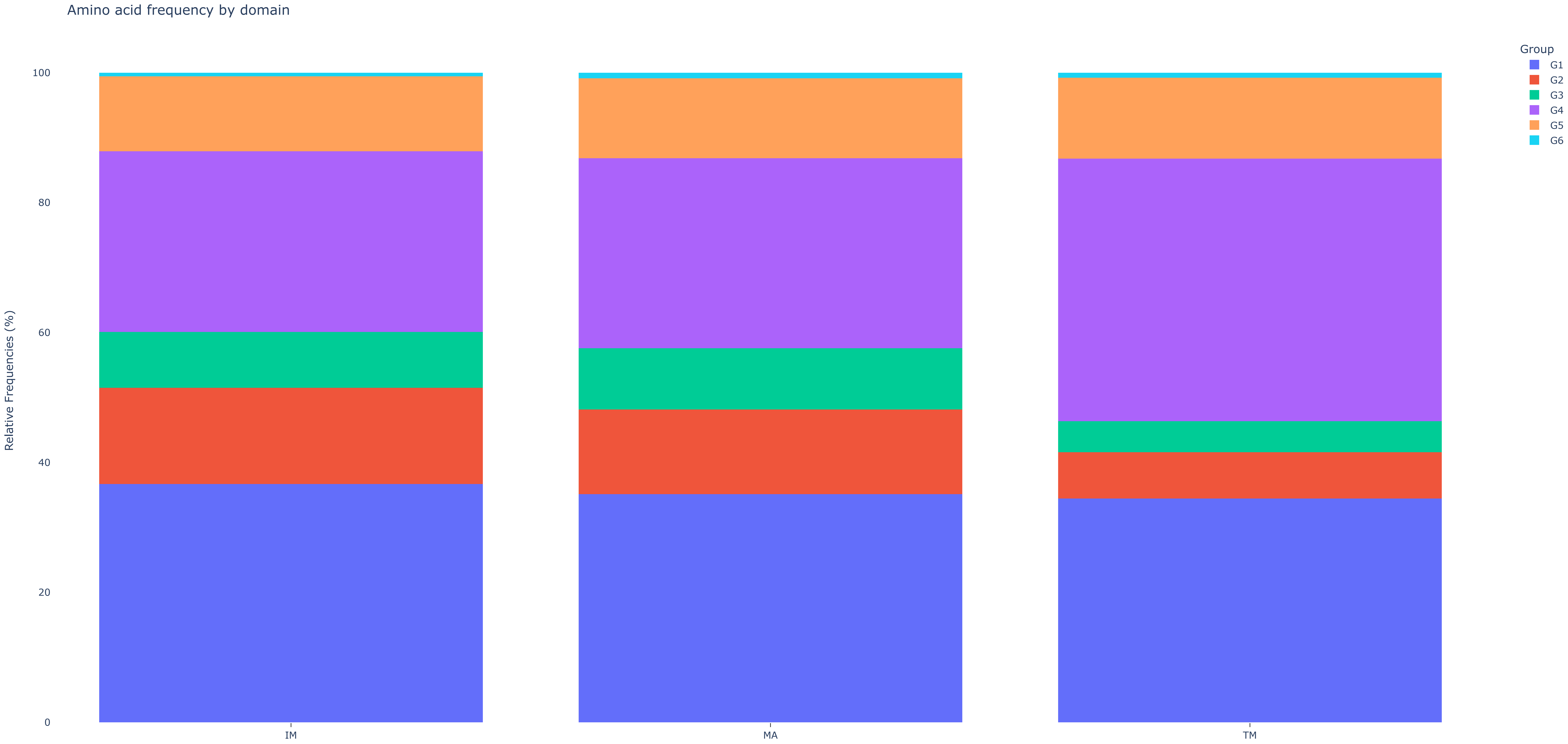


Loading [MathJax]/extensions/MathMenu.js
