## Supplementary material for "Integrating Secondary Structure Information Enhances Phylogenetic Signal in Mitochondrial Protein Coding Genes": Supplemetal TableS1

Table S1. AICc values associated with different partition schemes following merging/optimized with Partition Finder. AICc values are given for the first row only while the difference in percentage, relative to the first one, is given for the following rows. Colours indicate those partitions schemes that retrieved second codon sites of TM helices as independent partitions after optimization in Partition Finder: from positive strand only (orange), each strand as single independent sets (blue), and both strands merged in one (red).

| partition/species | Collembola | *Hyalella* | *Metacrangonyx* | *Pseudoniphargus* | Mammals | Primates | Vertebrates |
| --- | --- | --- | --- | --- | --- | --- | --- |
| 18p cod str dom | 538146.9205 | 205148.7652 | 273188.9300 | 329605.4692 | 1407540.4087 | 600410.7850 | 1736323.9654 |
| 12p cod str dom(TMvsMA+IM) | -0.0003 | 0.0000 | 0.0038 | 0.0000 | -0.0030 | 0.0085 | 0.0416 |
| 9p cod dom | 0.9946 | 0.1006 | 0.3255 | 1.1499 | 0.9644 | 0.7602 | 0.6631 |
| 6p cod str | 0.0000 | 0.0752 | 0.1443 | 0.0601 | 0.0635 | 0.0814 | 0.0466 |
| 3p cod | 0.9946 | 0.1766 | 0.4746 | 1.2439 | 1.0952 | 0.8336 | 0.7558 |
| single | 3.5599 | 3.4235 | 3.9152 | 4.1525 | 4.2372 | 3.7205 | 3.4406 |

codon (cod), strand (str), transmembrane domain (dom)
